# Identification of Distinct Topological Structures From High-Dimensional Data

**DOI:** 10.1101/2025.08.26.672446

**Authors:** Bingxian Xu, Rosemary Braun

## Abstract

Single-cell RNA sequencing allows the direct measurement of the expression of tens of thousands of genes, providing an unprecedented view of the transcriptomic state of a cell. Within each cell, different biological processes such as differentiation or cell cycle take place simultaneously, each contributing a different characterization of cell state. To identify gene sets that govern these processes for the purpose of disentangling convolved biological processes, we present “Identification of Distinct topological structures” (ID). ID works by constructing an alternative low-dimensional parametrization of the high-dimensional system, applying a finite perturbation to this alternative parametrization, and looking for genes that respond similarly. With this approach, we demonstrate that ID is capable of identifying structures within the data that will otherwise be missed. We further demonstrate the utility of ID in scRNA-seq datasets collected under various conditions, delineating cellular differentiation, characterizing cellular response to external perturbation, and dissecting the effect of genetic knock-outs.

## 1 Introduction

Dynamical processes such as differentiation [1, 2, 3, 4], cell cycles [5, 6, 7, 8], and circadian rhythms [9, 10, 11, 12, 13] are fundamental to life. Together, these processes define cellular state in a multi-faceted way. Single cell transcriptomic profiling (scRNA-seq) now provides a powerful means to investigate the interplay of these processes by characterizing the gene expression patterns associated with these processes. Because of the high dimensionality of modern transcriptomic assays, scRNA-seq analysis typically begins with feature selection, often based on signal-to-noise characteristics, to define a reduced set of informative genes. Cell–cell distances are then computed in this restricted feature space and embedded into lower dimensions using methods such as Uniform Manifold Approximation and Projection (UMAP) [14], enabling visualization of the data’s global and local structure [15, 16, 17, 18, 19].

However, different biological processes will induce different topological structures in the gene expression space. For example, cell cycle dynamics are intrinsically periodic and should therefore manifest as a ring-like topology, whereas differentiation trajectories are hierarchical and more naturally represented as tree-like structures. These will in turn give rise to different notions of “distance”; for instance, two cells may be proximal with respect to lineage identity, yet distant in cell cycle phase (or vice versa). As a result, distances computed from a generic feature set may conflate distinct processes and fail to accurately represent any one of them. The goal of this work is to identify the genes that define these different biological processes and use them to unravel the topology of the data.

This problem is traditionally solved by some form of matrix factorization, where the original gene expression matrix is decomposed into simpler components. Constraints imposed on the factorization can enhance its interpretability. For example, by requiring the factorized matrix to be non-negative using non-negative matrix factorization (NMF) [20, 21, 22, 23, 24], the gene expression of each cell can be viewed as the sum of the different gene programs it activates. However, gene programs identified by NMF can still appear as arbitrary collections of genes rather than biologically coherent sets [25, 26, 27]. Additional constraints can instead be imposed by using prior knowledge. For example, Spectra [27] uses user-defined gene sets to guide the factorization of the gene expression matrix. Similarly, CellUntangler [28] uses a variational auto-encoder to construct a latent space that best captures the topology of a user-defined gene set.

In real-world settings, the best–known marker genes of a biological process may not have been detected in the data due to low read depth, which limits the applicability of the aforementioned methods. However, as these markers often drive the regulation of other genes, the data can retain signatures of the relevant topology even as the marker genes themselves remain undetected. Following this logic, scPrisma [29] exploits prior information about the topology of the data (e.g., the existence of periodic or linear signals) by conducting spectral template matching between the spectrum of the data and the theoretical covariate spectrum expected from a user-defined topology.

Still, especially during the data exploration phase, neither the topology nor the relevant gene programs may known ahead of time. Because of this, an *unsupervised* algorithm is needed to identify gene sets that define the various aspects of the data. GeneTrajectory [30] is built for this purpose. Through the construction of a gene-gene distance matrix via optimal transport on a cell-cell graph, GeneTrajectory constructs an embedding for genes that allows direct visualization of the topological structures present in the data and identifies genes responsible for each structure. However, the construction of a cell-cell graph implies that the data contains only a single underlying biological process. For example, consider a population of cells that is both undergoing division (cell cycle) and differentiation; the cell-cell similarity graphs defined by the cell cycle genes, the differentiation genes, or the union of all genes may be different. In this situation, methods that rely on a single cell-cell graph, such as GeneTrajectory [30], may obscure biologically relevant relationships between cells or classification of genes; instead, it may be more informative to define the relationship between cells using multiple cell-cell graphs that reflect different processes.

To address the difficulty of conducting unsupervised identification of gene sets that define different topological structures, we developed *Identification of Distinct topological structures (ID)*, which takes only the count matrix as input to generate an embedding for genes with which they can be clustered. ID achieves state-of-the-art performance by first constructing an alternative low–dimensional representation of the system and then identifying genes that respond similarly when the alternative parametrization is perturbed. The intuition behind ID is that genes that are related to the same biological process should respond in concert to the same perturbation. We tested the performance of ID in multiple synthetic datasets and evaluated how it compares to existing unsupervised approaches. We then applied ID to multiple scRNA-seq datasets to unravel structures hidden within the data, and uncover how cells respond to both environmental and genetic perturbation. Lastly, using two independent human lung datasets, we “IDed” a conserved gene set that robustly defines lung epithelial differentiation.

## 2 Results

The state of a cell can be defined by the genes it expresses, and the distance between two cells may be defined by the difference between their states in the transcriptomic space. These distances can be used to visualize the cell state manifold and deduce cellular trajectories [31, 17, 32]. However, if the distances defined by different subsets of genes disagree with one another, the cell state manifold they construct would also differ; what, then, is the “correct” cell state manifold, and how can we identify sets of genes that contribute to different interpretations of the data?

By way of example, Figure 1A illustrates data occupying a 3D torus. When the data are projected to the space defined by gene 1 and gene 3, a linear structure (from light to dark blue) is apparent, but the cyclic pattern is lost. Similarly, in the case of scRNA-seq data, some genes may jointly define a linear trajectory (e.g., the aging of cells) while others may define branched (differentiation) or cyclic (cell cycle, circadian) trajectories. In these separate spaces, notions of cell–cell similarity (and hence inferences based upon them) may be very different.

**Figure 1.**
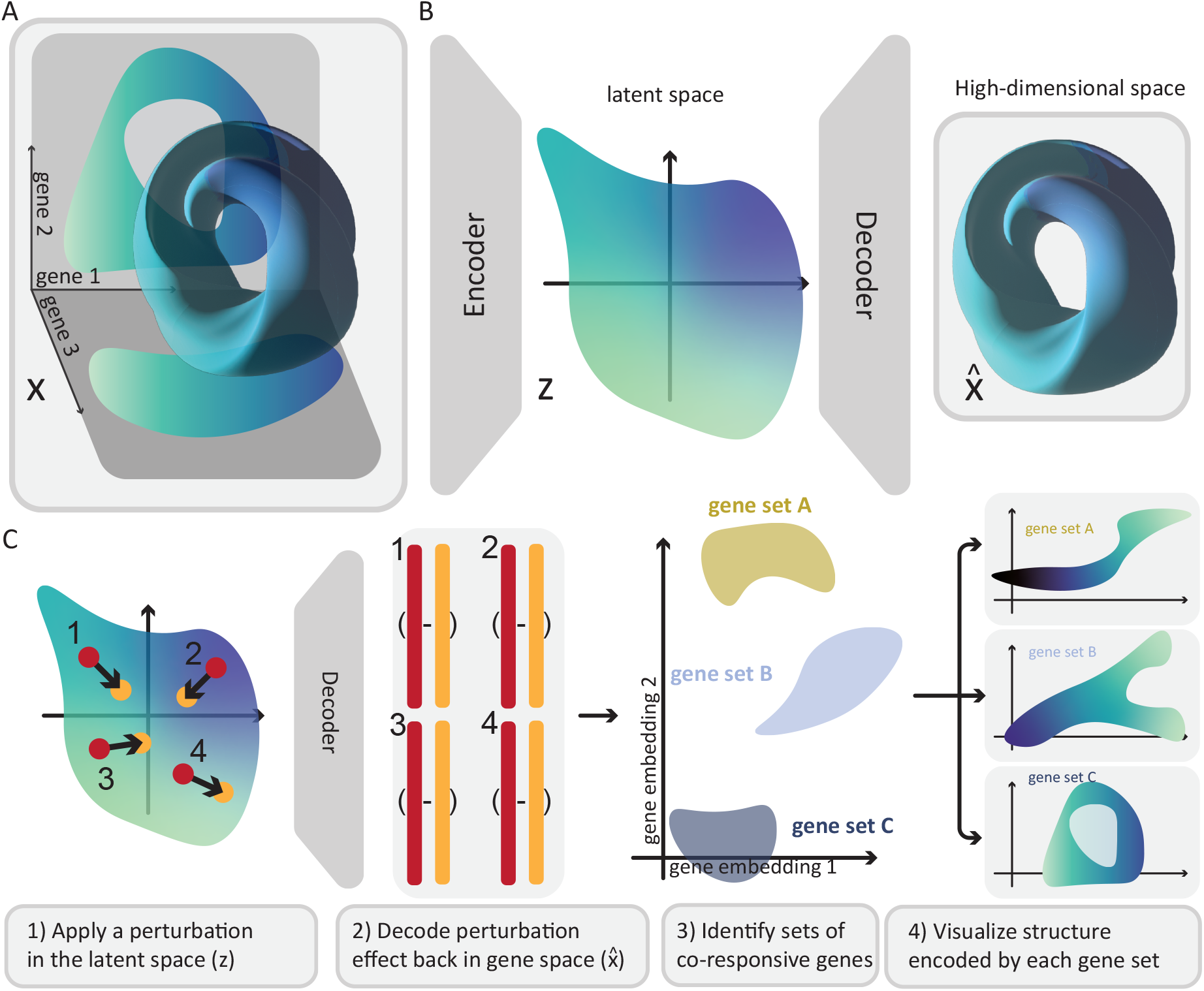
ID overview. A: Example of a 3-dimensional system with a cyclic and linear component. The gene 1 – gene 2 space exhibits a cycle, whereas and the gene 1 – gene 3 space describes a linear trajectory from light to dark blue. B: In ID, high dimensional data (*x*) is mapped onto a low dimensional space (*z*) via a variational autoencoder. C: The latent representation of the cell is then perturbed in different ways, and the response in the original space 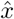 (output by the decoder) is used to identify different gene sets by clustering their response.

To identify gene sets that contribute to various topological structures, and hence different biological processes and measures of cell similarity, we developed *Identification of Distinct topological structures (ID)*. The key output of ID is an embedding of genes, wherein each cluster contains genes that define similar cell–cell distances.

Our method for identifying feature sets that define particular topologies in the data begins from the assumption that living cells lie on a low–dimensional manifold within the high–dimensional gene expression space [33, 34, 35, 36, 37, 38, 39]. Learning this low–dimensional representation allows us to study how changes on the low–dimensional manifold are reflected in the full gene expression space. We postulate that genes that jointly define a “coherent” structure will respond similarly when a perturbation is applied in the low–dimensional representation. With this reasoning, we hypothesized that if we first find a parametrization of a high-dimensional system (*x*) that maps it to a lower-dimensional space (*z*) (Figure 1B), we may be able to cluster features based on how they respond to a small perturbation of the latent representation of the original data, and that each cluster of genes will jointly define a unique topological structure (Figure 1C).

### 2.1 The ID algorithm

ID begins by learning a mapping between the high–dimensional gene expression space and a low– dimensional embedding. Given a raw data matrix, we first standardize the mean and variance of each feature by *Z*-scoring across samples to obtain the data matrix we use as input, **X** ∈ ℝ^*N* ×*S*^, where *N* and *S* are the number of features (genes) and samples respectively. For scRNA-seq count matrices, we normalize by the library size of the cell before *Z*-scoring each feature. We then train a variational autoencoder (VAE) [40] to map ***x***_*s*_ (the data vector for sample *s*) to a low–dimensional latent space ***z***_*s*_ and then back to 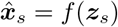 (Figure 1A,B), where *f* (·) is the decoder network learned by the VAE. Optimization of the VAE is described further below.

We then select a random point ***z***_*j*_ in the latent space, apply a perturbation *δ****z***_*j*_ (Figure 1C, step 1), and decode the perturbation back into the gene expression space to obtain a perturbed 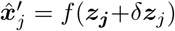. We have set the perturbation magnitude to |*δ****z***| = 0.1 by default. From this we construct a response matrix, **M** ∈ ℝ^*N* ×*P*^, where *P* is the number of computational perturbations *j* that we perform. Each entry of **M**, *m*_*ij*_, denotes how gene *i* responded to the *j*^th^ perturbation, and is the key to our identification of related features. The *j*^th^ column of **M**, denoted by ***m***_*·j*_ is computed by:

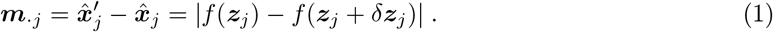

It is easy to see that the *rows* of **M, *m***_*i·*_, represent how each gene *i* responded to the *P* minor perturbations in the latent space. By sampling multiple points randomly in the latent space as well as random directions, and plugging them into Eq 1, we assemble the response matrix **M**. Finally, we scale each column of **M** via *Z*-scoring to obtain the scaled response matrix, 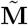.

By default, we set the number of perturbation trials, *P*, to 50, 000. To obtain sets of co–responsive genes (Figure 1C, step 3), we first use principal component analysis to extract the top 20 PCs of 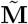 and subsequently cluster the *N* genes in a 2D UMAP space.

### 2.2 ID identifies distinct topological structures

To test how ID performs, we constructed a toy dataset that contains a linear and a branched structure (Figure 2A, first panel, and Figure S1B) that follows a “similar order” along both structures. In other words, as cell progresses along the linear structure, it also progresses somewhat smoothly along the branched structure. (An example where cells follow a different order can be found in Figure 2B.) We first defined the linear and the branched topology on the two-dimensional space and projected each structure into a 50-dimensional space, resulting in a gene expression matrix that contains 1000 cells (samples) and 100 genes (features) for each cell (Figure 2A, second panel). By analyzing the responses of the genes to perturbations using ID, we observed that genes contributing to the two structures form two well-defined clusters (Figure 2A, third panel). Alternatively, when we attempted the naïve approach of using UMAP [14] on the transpose of the gene expression matrix (effectively clustering the gene expression directly), we observed that this approach failed to separate the two gene clusters (Figure 2A, inset in the third panel). Thus in this simple test, ID at least out–performs the most naïve of approaches. We then sought to test how UMAP–based clustering would perform if the two structures were more “different”. To do so, rather than introducing another topological structure, we simply changed the order of cells along each structure such that the two structures were less correlated with one–another. In this example, when a cell travels along the “Y”-shaped structure, it no longer traverses the linear structure smoothly but instead jumps around (Figure 2B in contrast to Figure 2A). With orders of cells altered, we observed that the UMAP projection of the transpose of the gene expression matrix is able to produce two distinct clusters, each representing a distinct topological structure (Figure 2B). Thus, in cases where the gene sets define completely orthogonal trajectories, they can be clustered in the gene–UMAP space (Figure 2B), but in cases where there is some overlap — as might be expected in real data due to coordination of biological processes — ID is better able to articulate the relevant gene subsets (Figure 2A).

**Figure 2.**
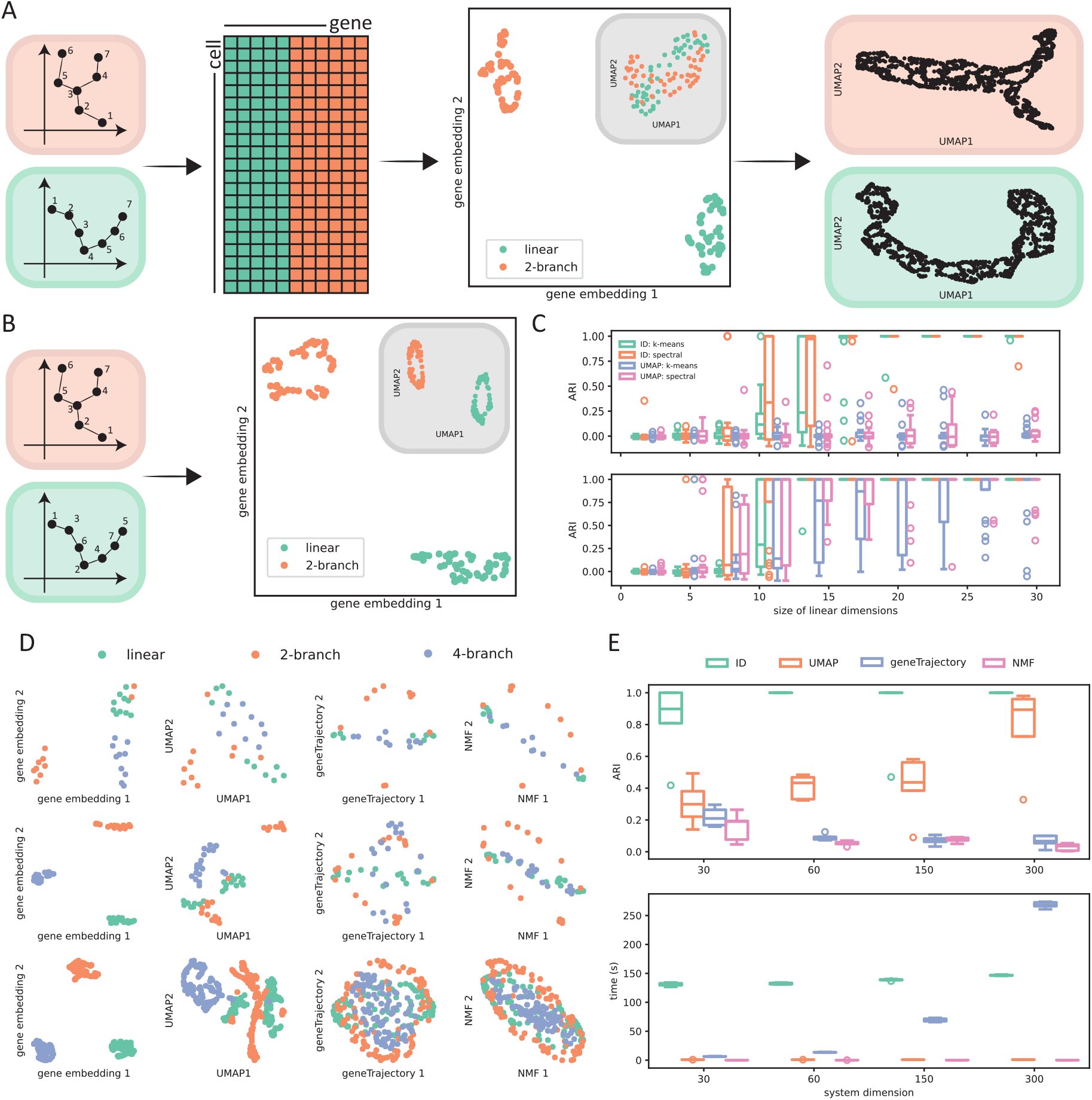
Applying ID to simple toy datasets. A: Illustration of how ID identifies sets of genes that define different structures in the data. We first constructed two topological structures on a low-dimensional space (panel 1) and mapped them both to a high-dimensional space of “genes” before concatenating them together (panel 2). We then used ID’s perturbation responses to construct a gene embedding (panel 3; for comparison, a simple UMAP embedding of the genes is shown in the inset, where it is apparent that the two sets of genes are not separable). We then constructed UMAP embeddings of the cells using the subsets of genes in the two ID clusters, and observed that the clusters indeed define a linear and a branched sub-component as expected (panel 4). B: An example where the “order” defined by the two topological structure differs greatly. In this case, both ID and gene-UMAP allow perfect identification of the genes contributing to the two structures. C: Comparing ID and UMAP in classifying genes when the number of features defining the linear component ranges from 2 to 98 (plot truncated at 30; the full figure may be found in Supplementary Figure S2). Top: examples when the “order” is similar. Bottom: examples when the “order” is different. 20 experiments were conducted for each condition. D: Example embeddings of genes produced by different algorithms in toy datasets that contain three different structures. The toy dataset used in first, second and third row contained 30, 60, 300 genes respectively. E: Accuracy as measured by the adjusted Rand index (ARI) and the run time for each method as a function of total system dimension. Each trial was repeated five times.

We next sought to investigate how separable these structures would be as a function of the number of genes defining each of them. To investigate how ID performs when the dimensionalities of the linear and the branched component differ, we fixed the system dimension to 100 and gradually increased the size of the linear dimension from 2 to 98. When the two topological structures share a similar ordering of cells (as in Figure 2A), we observed that ID failed to separate features when the size of the linear dimension was too small, having an adjusted Rand index (ARI) close to 0. As the size of the linear dimension increased above 15 (out of 100), ID started to have a near perfect performance (Figure 2C, top panel; Figure S2, top panel). In general, we observed that spectral clustering and *k*-means clustering performed similarly. In sections that follow, we used *k*-means clustering by default for its computational efficiency unless otherwise noted. As expected from the previous test, when the order defined by the two classes of features is similar, the UMAP projection of the transpose of the gene expression matrix cannot separate the two classes of features regardless of their ratio (Figure 2C, top panel; Figure S2, top panel). Similarly, when the order of cells along the two processes is different, we observed that neither ID nor UMAP can separate the features by the topological structure they represent when the size of the linear dimension is too small. However, consistent with previous observations, both approaches begin to have near-perfect performance when the size of the linear dimension grows larger than 15 (Figure 2C, bottom panel; Figure S2, bottom panel).

Next, we compared the performance of ID against two additional well–established methods: non-negative matrix factorization (NMF) [20, 21, 22, 23, 24] and geneTrajectory [30]. To test how each method performed, we first conducted the comparison in a situation with three distinct topological structures, each composed of the same number of genes. By changing the total dimension of the system, we observed that ID out-performed the other three methods (UMAP, NMF, geneTrajectory) in discerning constituents of the three topological structures regardless of the dimension of the system (Figure 2D). In this test case, NMF and geneTrajectory consistently under-performed while the performance of UMAP improved as the size of the system increased (Figure 2E, top panel). We also tested how the run time of each method scales with the number of features. We observed that when the size of the system is small, ID is the least efficient method, but as the size of the system increases, the runtime of geneTrajectory scales exponentially and rapidly exceeds that of ID, most likely as a consequence of its internal construction of a gene-gene distance matrix (Figure 2E, bottom panel). By contrast, the runtime of ID scales linearly with sample size; for typical scRNA-seq datasets with a few tens of thousands of cells, ID completes in within a few minutes on Intel Core i7-10700 (8-core) CPU.

In the previous example, when we used NMF, we decomposed the gene expression matrix into three components because we knew that our toy data contained three topological structures (see Methods for details). However, we observed that the performance of NMF appeared to depend non-linearly on the number of components that we decompose the data matrix into (Figure S3A). By decomposing the data matrix into a wider range of components and computing the resulting ARI, we observed that the performance of NMF increased when we decomposed the data matrix to 30 components but then decreased when we attempted to decompose our data further (Figure S3B). However, without knowing the structures (and thus which genes belong to them) ahead of time, one cannot use the ARI to find the optimal number of components to decompose the data matrix. To examine whether there are alternative approaches to select the optimal number of components, we conducted a much finer sweep (Figure S4). When we plotted the NMF reconstruction error against the number of components, we observed that the elbow region of this curve overlaps with the ARI peak. This thus provides a potential way to select the optimal number of components when using NMF. We also investigated whether similar problem is also present in ID by testing how it performs when we change the dimension of the latent space. Unlike NMF, we observed that ID can achieve a near perfect performance as long as the dimension of the latent space is sufficiently high (Figure S5); thus, in contrast to NMF, ID does not require a sweep (which would increase the computational cost of the NMF approach).

Next, we investigated how ID would perform in a more complex scenario. In this test case, we sampled from the linear and branched structure, as well as six other distinct topological structures (Figure 3A). In total, we sampled from eight different structures, and projected each onto a 50 dimensional space (from the original 1–3) via a topology preserving transformation (see Methods for details), yielding a final data matrix with a 400–dimensional feature space of “genes.” When we visualize this data on a 2D plane using UMAP, we observe that the low–dimensional projection of cells has a branched structure but displays no characteristics of the 3D shapes described by the other “genes” (Figure 3B). Interestingly, when we swept across the number of components for NMF, we observed that the peak position has shifted from our previous example (Figure S6A and Figure S7), reflecting the greater complexity. However, as noted above, because the run time for NMF increases as the number of components increases, the need to sweep and search for the optimal number of components makes NMF much more computationally expensive than ID (Figure S7, bottom panel). In addition to ID’s efficiency, we observed again that ID can separate the distinct classes of features well, out-performing all other methods and achieving near–perfect performance in almost all trials (Figure 3C, D and Figure S6B).

**Figure 3.**
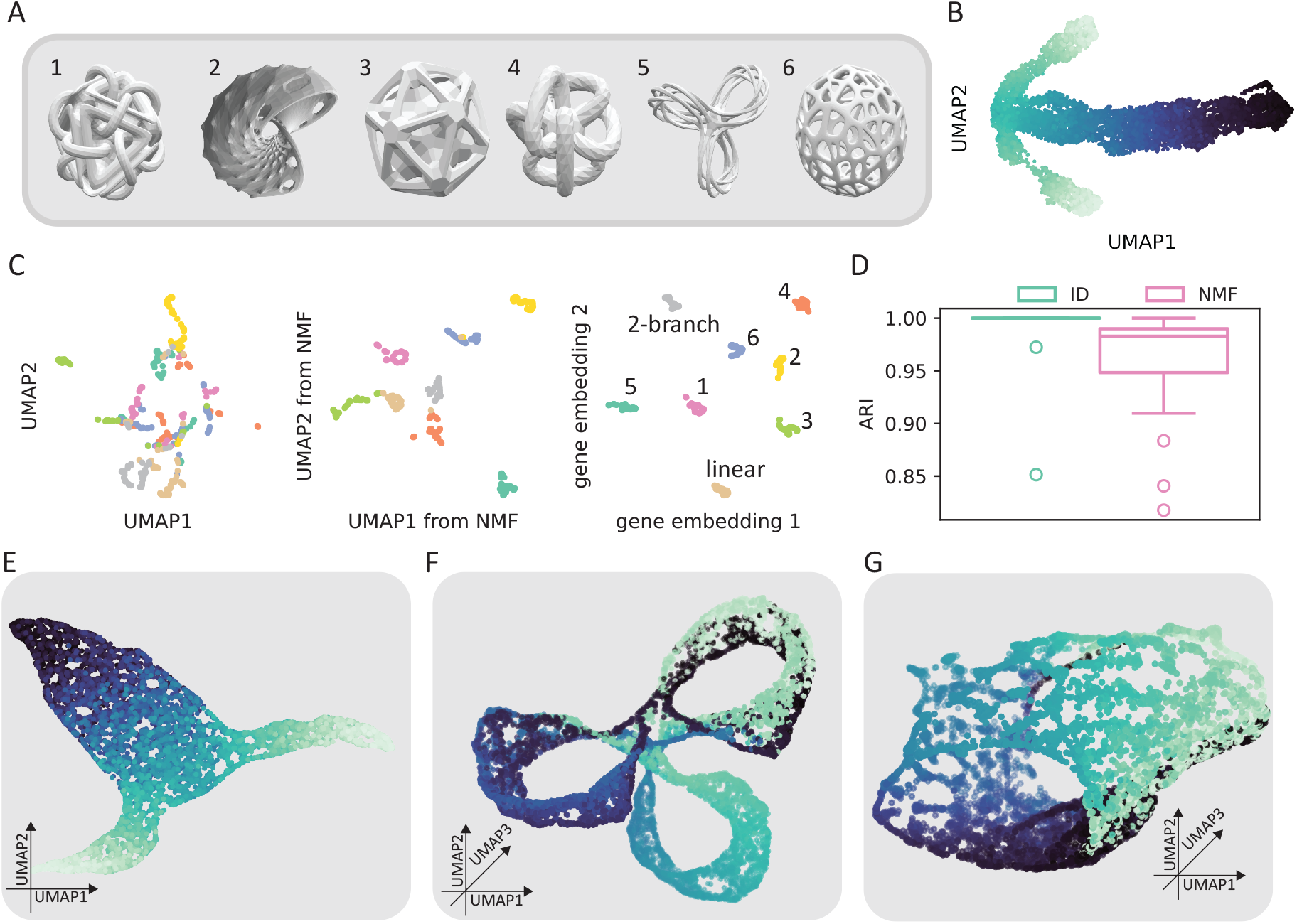
Applying ID to a complex toy dataset. A: Manifolds from which we sampled to construct the toy dataset, in addtion to the linear and branched structures previously described. 1, Octahedron; 2, Clifford torus; 3, Dodecahedron; 4, Genus-6 torus; 5, Genus-3 surface; 6, Voronoi sphere. B: UMAP visualization of samples constructed using all features; only the branched structure is evident. C: Embedding for genes constructed by UMAP, NMF and ID; ID clusters are well–defined and correspond to the structures present in the data. D: ARI computed using k-means clustering on the embedding for genes constructed from NMF or ID from 20 individual trials. E-G: UMAP projections of cells using only the grey (E, 2-branch), dark green (F, cluster 5: Genus-3 surface) and yellow (G, cluster 2: Clifford torus) from Figure 3C.

For each cluster of genes we IDed, when we visualize the data with UMAP, we observed that the UMAP–embedded data reflects to low-dimensional topologies with which we constructed the high-dimensional data (Figure 3E-G and Figure S6C). Together, our results show that ID can reveal distinct topological structures that are masked within the data.

### 2.3 ID delineates cellular differentiation

To illustrate how ID can disentangle and elucidate biological processes, we applied it to several scRNA-seq datasets. We first considered two datasets that respectively charted the differentiation of hematopoetic stem cells [41, 42] and the development of the mouse hippocampus [43, 44]. Applying ID to these two datasets revealed two major clusters of genes in their gene embeddings: a larger cluster of genes that define a tree-like topology, corresponding to the annotated cell types; and a smaller cluster of genes that define a ring-like topology, corresponding to the phases of cell cycle (Figure 4A, B). We also noticed the presence of small clusters of genes that are very tight and distinct from the main clusters (such as the orange cluster in Figure 4B); these typically comprise genes with low (or predominantly 0) expression, and thus contain little information. As illustrated in the first grey panels of Figure 4(A,B), the cluster-1 genes delineate the differentiation landscape of these cells.

**Figure 4.**
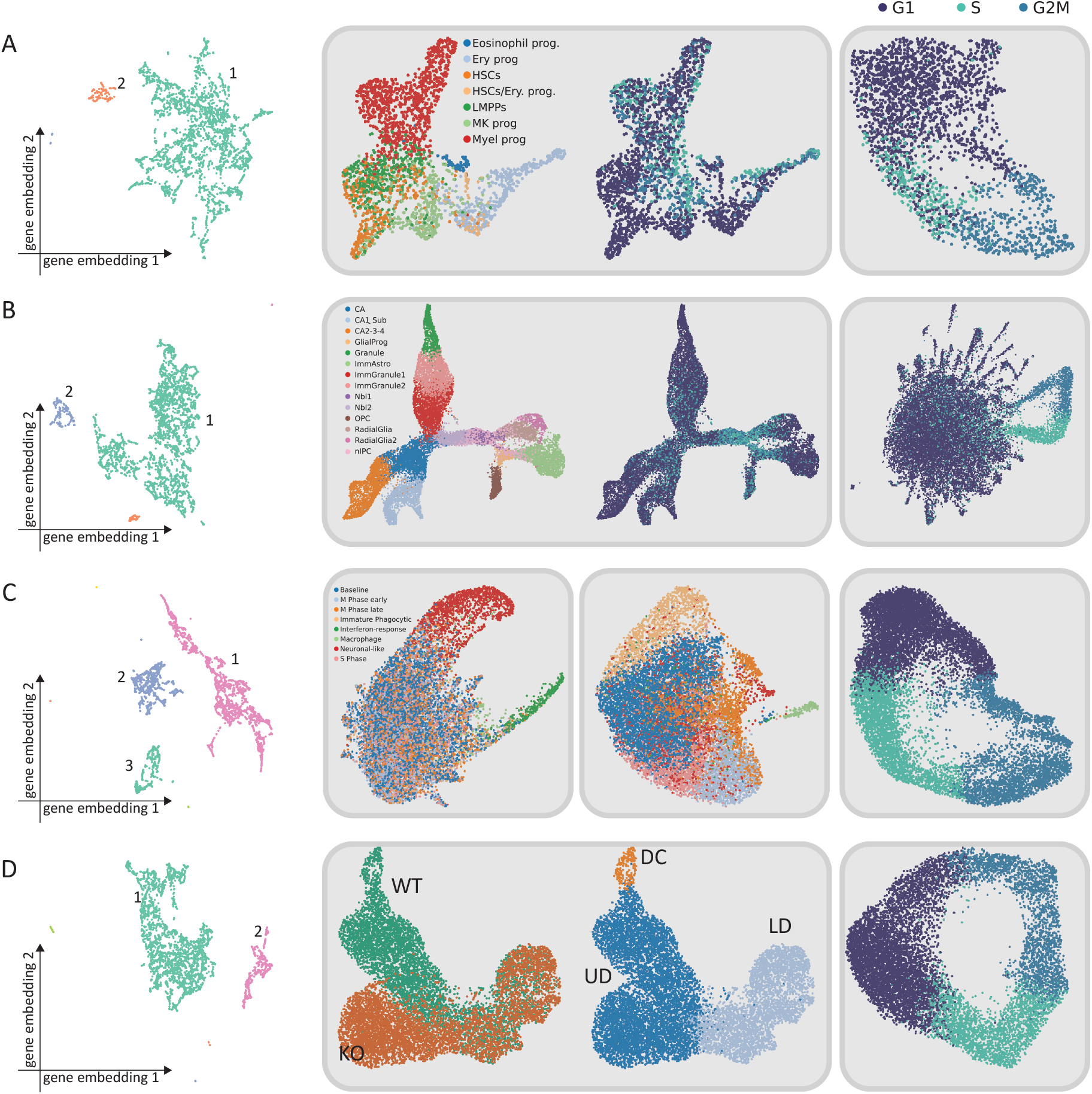
Applying ID to scRNA-seq datasets. ID is applied to (A) hematopoetic differentiation [42, 41], (B) hippocampus development [43, 44], (C) microglia [45], and (D) hair follicle [30] scRNA-seq datasets. The resulting gene embedding and gene clusters obtained for each dataset is plotted on the first column. In each row, each box contains the UMAP projection of cells constructed using the respective gene clusters (e.g., the UMAP in the first box is constructed from the cluster 1 genes, etc.)

Given the large size of cluster 1 in both cases, one may reasonably ask whether one would obtain similar inferences simply using all genes. To investigate if it is worthwhile to analyze the data with just the cluster-1 genes (which describe the tree–like structure), we produced a UMAP embedding of cells using all features (Figure S8A, B) and compared them to those created with only the cluster-1 genes (Figure 4A, B). While we observed that these embeddings are largely similar, the UMAP generated from all genes appeared confounded by other processes. For example, in the hematopoetic differentiation dataset, we observed that a group of lymphoid-primed multipotent progenitor cells (LMPPs), megakaryocyte (MK) progenitors and HSCs, protrude between the branch of erythroid progenitor and myeloid progenitor, almost forming an additional branch (Figure S8A) despite not being a different cell type. Closer examination showed that these cells are in either the G2 or the S phase of the cell cycle (Figure S8A). When only the cluster-1 genes were used, this artefactual branch disappeared (Figure 4A). Similar phenomena were also observed in the hippocampus differentiation dataset; when all genes were used for the UMAP, the neural intermediate progenitor cells (NIPCs) and oligodendrocyte progenitor cells (OPCs) also appeared to have additional branches (Figure S8B) that were driven by the cell cycle (Figure S8B), whereas if only the cluster-1 genes were used, these branches disappeared (Figure 4B). Interestingly, in the new embedding, we find that the NIPC did not even turn into its own segment, but rather merged into the main branch coming from Nbl1 cells and extended to both the radial glia and the glial progenitors (Figure 4B). This observation suggests that this NIPC population is either mischaracterized as a unique cell type or is defined by persistent cell cycle activity. If the latter is true, it raises the question of whether the cell cycle component should be regressed out during single cell analysis.

To sum up, using two differentiation datasets, we illustrated that ID is capable of identifying gene sets that define distinct biological processes. It may be important to use these gene sets separately, as our examples demonstrate that their joint usage can introduce additional branches in the differentiation tree, affecting downstream analyses such as trajectory inference.

### 2.4 ID reveals cellular response to external perturbation and articulates discrete state transitions

We next demonstrate how ID can reveal biologically relevant information with an scRNA-seq dataset of microglia [45]. Microglia are macrophages that reside in the brain and play an important role in the development of the neuronal system, with diverse transcriptional states that are associated with different functions and morphologies. Here, we examined scRNA-seq data of microglia sampled from mice with or without whisker deprivation [45]

When all genes were used to construct a UMAP representation, we obtained a projection consistent with the original annotation, which is a combination of the cell cycle phase and annotated cell types (Figure S8C). Notably, while all phases of the cell cycle were well represented, this low-dimensional projection contained no obvious circular structure (Figure S8C).

By applying ID, we identified three major clusters of genes (Figure 4C). Interestingly, we observed that the largest cluster of genes only discriminates the neuronal–like and the interferon–responsive microglia, while a second, much smaller cluster of genes defines the majority of the annotated clusters (Figure 4C). We also observed that the third cluster of genes is associated with the cell cycle, evident from the low-dimensional projection of the cells constructed using these genes, which both formed a clear circular structure and is consistent with the cell cycle phase estimation (Figure 4C).

The original study suggested that whisker removal would lead to the expansion of an interferon– responsive microglia population which are capable of engulfing neurons. This would limit the accumulation of DNA–damaged cells and generate a population of microglia that express engulfed neuronal markers [45]. The relationship between the interferon–responsive microglia and neuronal–like microglia was previously posited via RNA-velocity [43]. However, if this group of interferon–responsive microglia start to “express” neuronal markers due to their engulfment of DNA-damaged neurons, how could RNA velocity, which attempts to predict the future transcriptome of a cell, anticipate the cellular target of microglial phagocytosis? This question led to our closer examination of the low-dimensional projection of cells constructed from the cluster 1 genes, which can discriminate these two cell types. Interestingly, we observed that this low-dimensional projection contained a “gap” between the interferon–responsive and the neuronal–like microglia that is bridged by other microglia populations (Figure 4). This is consistent with the notion that the transition between the transcriptional state of a microglia with and without an engulfed neuron should be discrete. In order to engulf other cells, neuronal–like microglia needs to break down the engulfed neuron and return to a “ground-state” cell before transitioning to an interferon-responsive cell [46, 47, 48]. That is, with reference to the first grey panel in Figure 4: a green interferon–responsive microglia engulfs a neuron, making a discrete leap to the red neuronal–like microglia; it then digests the neuron and returns to the interferon–responsive green state via the other (mixed) microglia states.

To summarize, we observed that this microglia dataset contained multiple biological processes which can make interpretation of the data challenging when they are not deconvolved (Figure S8C). By using ID, we identified three distinct topological structures from the data, corresponding to biologically relevant processes [45]: the cell cycle, cell identity, and engulfment of neurons by microglia.

### 2.5 ID dissects the effect of gene knock-outs

Hair follicle dermal condensates are essential for the formation of hair follicles, whose differentiation is regulated by the Wnt signaling pathway [49, 50, 51]. Here, we examined scRNA-seq data of dermal cell differentiation sampled from wild-type and Wnt knock-out mice [30]. This defect is known to result in the lack of formation of dermal condensates, and consequently the lack of hair follicles [30, 52, 53].

When all feature genes are used, the UMAP projection describes cellular identity well, clearly separating the lower dermis (LD), upper dermis (UD) and dermal condensates (DC) as shown in Figure S8D (center panel). However, it fails to segregate cells by genotype (Figure S8D, left), and an apparent branch in the structure is dominated by cell cycle (Figure S8D, right).

By applying ID, we observed two major clusters of genes (Figure 4D). Dimensionality reduction using cluster 1 genes as features revealed a clear, linear structure, where the lower dermis is more similar to the upper dermis than to the dermal condensate (Figure 4D). More importantly, the effect of Wnt knock-out is directly visible along this structure (Figure 4D). On this space, we observed that while the lower dermis of the WT and KO mice are transcriptionally similar, that of their upper dermis is not, suggesting that Wnt has disrupted the formation of dermal condensates by altering the transcriptional state of the upper dermis. As in the previous examples, we have also identified a cluster of genes that define the cell cycle. The low-dimensional projection of cells defined by these genes form a clear circular structure that is consistent with cell cycle phases (Figure 4D).

### 2.6 ID identifies conserved differentiation trajectory in human lungs

So far, we have demonstrated how ID can delineate cellular differentiation (Figure 4A, B), elucidate the cellular response to external perturbation (Figure 4C), and dissect the effect of gene knock-outs (Figure 4D) by identifying genes that define distinctive topological structures. Next, we use ID to test the hypothesis that, for a given system, there will be a conserved set of genes that are responsible for defining common topological structures. In other words, we seek to find datasets that characterize the same system, apply ID to these datasets, and cluster genes accordingly to investigate if the cluster assignment of genes are consistent across datasets. As different datasets may be collected by different research groups or processed with different protocols under different conditions, they will inevitably contain systematic differences. Some of these differences may be attributable to changes of the transcriptional state of the cell incurred by the environment, and others may simply be gene expression noise or batch effects. By examining whether the ID-identified clusters of genes are consistent across datasets, we may assess the extent to which ID identifies genes associated with a biological process, as opposed to noise or technical artifacts. To this end, we analyzed two scRNA-seq datasets collected from the human lungs, hereafter referred to as the lung sample data [54], collected directly from two unsuitable transplant lungs; and the lung organoid data [55], collected from two donors which were then cultured into different organoid models.

We first examined the lung organoid data [55]. When all features were used to construct a low-dimensional representation of cells, we observed that each organoid model occupies a distinct region of the space (Figure S9A) and the annotated cell types can be well separated within each such region (Figure S9A). Similar to our previous examples, we also observed on this space the formation of clusters of cells consequent to their cell cycle phases (Figure S9A). Interestingly, we observed that cells sampled from the EX (PneumaCult EX-plus medium) model had a Y-shaped path of differentiation. Closer examination of the metadata revealed that each branch of the “Y” is composed of cells that were harvested from the same donor (Figure S9A, inset). In summary, when all features were used to construct a low-dimensional representation of cells, the resulting space is jointly affected by cell type, cell cycle phases, organoid model, and donor from which the cells were harvested.

Applying ID to the lung organoid data, we observed three major clusters of genes (Figure 5A, inset), with the largest cluster defining a space for cells that is similar to that constructed from all features. We then explored the low-dimensional space defined by the other clusters of genes. On the space defined by the cluster 2 genes, we observed a conserved “barrel”-shaped differentiation trajectory (Figure 5B). Despite the differences among these organoid models, we observed that the O1 and O2 organoid had an almost identical barrel-shaped trajectory while that for the other two models, ALIEX (cells cultured in air-liquid interface with PneumaCult-ALI medium) and EX (cells cultured in PneumaCult EX-plus medium), showed considerable differences (Figure 5B). A similar observation was also made in the space defined by the cluster 3 genes, which was observed to be associated with the cell cycle. In this space, we observed a striking similarity among the low-dimensional projection of the O1, O2 and the EX model, all possessing a ring-like structure that is consistent with the annotated cell cycle phases (Figure 5B). Interestingly, the ALIEX model stood out from the other three models. In the space defined by the cluster 2 genes, the basal cells and secretory cells of the ALIEX model clearly differed from those of the other organoid models; and in the space defined by the cluster 3 genes, the ALIEX model is the only organoid model from which we failed to observe a ring structure (Figure 5B). While some of the organoid models remain distinct from each other in both the “differentiation” and the “cell cycle” space, the extent of their differences are much smaller compared to that illustrated in Figure S9A. In addition, we observe in both the “differentiation” and “cell cycle” space that the effect of the donor is almost completely absent in all four organoid models (Figure S9B), suggesting that ID may have identified a conserved set of genes that define the differentiation trajectory and cell cycle phases of lung epithelial cells.

**Figure 5.**
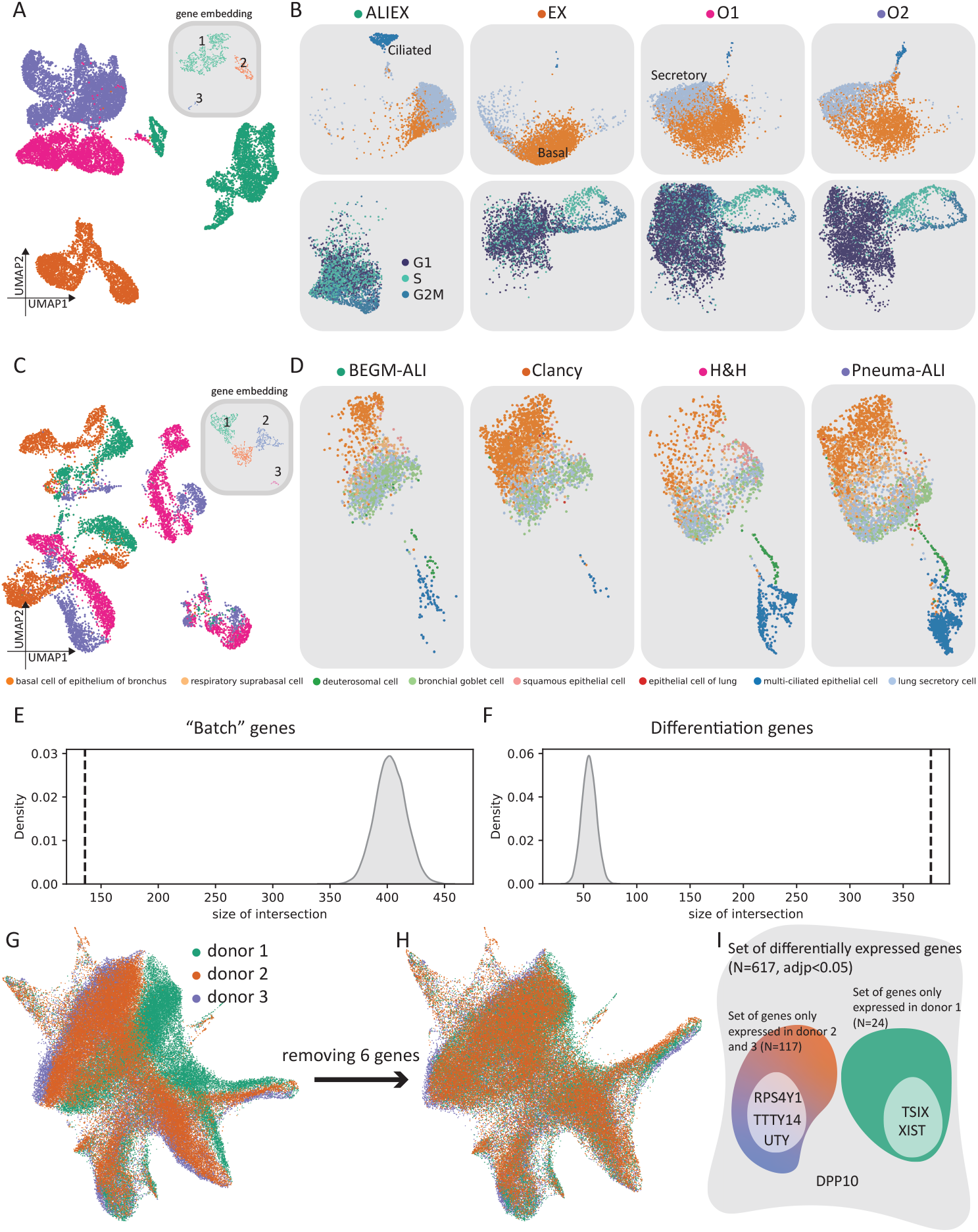
Applying ID to scRNA-seq datasets collected from the human lungs. ID is applied to the lung organoid dataset [55] (A, B) and the lung sample dataset [54] (C, D). A: UMAP projection of cells in the lung organoid data constructed from cluster 1 genes (gene embedding as inset). B: UMAP projection of cells constructed from cluster 2 (top) and cluster 3 (bottom) genes respectively. Each column contains cells in the same organoid model, and the same coordinate system is used for all columns. Colors denote cell type annotation. C: UMAP projection of cells in the lung sample data constructed from cluster 1 genes (gene embedding as inset). D: UMAP projection of cells constructed from cluster 2. Each column contains cells cultured in the same media, and the same coordinate system is used for all columns. Colors denote cell type annotation. E: The observed intersection between the cluster 1 genes from both datasets (dashed line) and the null distribution constructed by random sampling. F: The observed intersection between the cluster 2 genes from both datasets (dashed line) and the null distribution constructed by random sampling. G: Low dimensional projection of the HSC data, colored by donor. H: Low dimensional projection of the HSC data with 6 IDed genes removed. I: Summary of the identified differentially expressed genes.

These results suggest that while the lung organoid dataset contains a large amount of technical variation due to both the donor ID and the organoid model, ID can identify gene sets that are less influenced by this source of variation. This suggests that we have found a gene set that robustly defines the differentiation trajectory of lung epithelial cells, that is, it reflects a biological process rather than technical artifacts.

To validate this, we examined the lung sample dataset [54], which is derived from two discarded transplant lungs and subsequently cultured in four different types of media. Similar to the lung organoid data, the lung sample data also contains a large amount of technical variation, evident from the low-dimensional space constructed from all feature genes (Figure S10A). By using ID, we again identified multiple gene sets, with cluster 1 genes capturing most of the technical variations (Figure 5C). Strikingly, when we examined the space defined by the cluster 2 genes from the lung sample data, we again observed a “barrel”-shaped differentiation trajectory, which is conserved across both donors and culture media (Figure 5D and Figure S10B).

We then assessed the extent of overlap in the ID gene clusters. If the genes that defined the “barrel”-shaped differentiation trajectory are the same across the two datasets, then it would suggest that they are the defining properties of lung epithelial cell differentiation. To quantify the extent of the randomness of these gene sets, we tested, for a given gene set, if the observed number of intersecting genes would be different than if we were to sample genes randomly (see Methods for more detail). We first conducted this test for the cluster 1 and cluster 2 genes, which respectively captured the variation induced by experimental covariates and cellular differentiation. Interestingly, for the cluster 1 genes (which describe technical /batch variations) we observed that the number of genes that are conserved across the two datasets is significantly *smaller* than expected by chance; they are largely disjoint (Figure 5E). By contrast, the “barrel”–defining differentiation genes in cluster 2 showed a much greater overlap than expected by chance (Figure 5F). With the cluster 3 genes, we also observed that the number of intersecting gene to be much larger than the null model (Figure S10C). When we examined the low-dimensional space defined by the cluster 3 genes, we observed a ring-like structure similar to that observed in the lung organoid data (Figure 5B and Figure S10D).

To summarize, we first used ID to independently identify gene sets that defined cellular differentiation, cell cycle progression, and captured the most amount of technical variation in two human lung datasets. In both datasets, we observed that the variation introduced by experimental covariates is greatly reduced in both the differentiation and cell cycle space. To elucidate if the differentiation and cell cycle related genes that we discovered are biologically meaningful, we investigated if their identities are conserved across datasets. Via our permutation test, we observed both the differentiation and cell cycle gene sets to be highly conserved, while the gene set that captured the technical variation was highly different. This observation suggests that ID can be used to robustly and reproducibly identify key genes that define differentiation/cell cycle, and more importantly, that the expression of these key regulators is independent of factors such as donor or cell culture type.

### 2.7 ID detects and removes batch effects

We further explored the potential usage of ID on a dataset charting the differentiation of hematopoetic stem cells (HSCs) [56]. Using the conventional pipeline, we observed that the low-dimensional projection of the data varies across donors (Figure 5G). However, by applying ID, we identified a set of 6 genes (Figure S11A) whose absence removes this observed “batch”/donor effect (Figure 5H). Interestingly, we found these genes to be almost exclusively sex related, suggesting that the batch effect we observed is consequent to the sex of the donors. As expected, their removal has no effect on the structure of the cellular differentiation manifold (Figure S11B). To investigate if we can identify these six genes via alternative approaches, we conducted differential expression analysis and identified 617 genes showing differential expression, 117 of which are only expressed in male donors (donor 2 and donor 3), and 24 of which only expressed in the female donor(donor 1) (Figure 5I). By identifying a smaller, key set of genes with ID, corrections can be made without impacting other biologically relevant signal.

Together, this example demonstrates that ID can be used to identify “batch” drivers. In the above example, we showed that the removal of 6 genes completely removes what appears to be a batch effect. If batch correction methods were used, they would transform all genes, altering gene-gene relationships and potentially changing the result of downstream analysis. With ID, we are able to identify the source of the batch effect while leaving all other genes untouched.

## 3 Conclusion

We have introduced ID, a powerful and efficient algorithm to uncover hidden topological structures from data using computational perturbations. We demonstrated the utility of this approach through extensive benchmarks on toy datasets and comparisons to other existing methods. Our results demonstrate that when the state of the cell is jointly influenced by multiple extrinsic and intrinsic factors, ID is capable of identifying gene sets associated with distinct topological structures and dynamical processes.

Through our investigation, we observed that distinct gene sets can provide differing measures of cell-cell distance. For example, cells of the same cell type may not be in the same phase of the cell cycle. As a result, the joint usage of cell cycle genes and cell type markers to define cell state can confound cell type annotation and produce misleading low-dimensional visualization of the cells or incorrect trajectory inference (Figure 4C, D and Figure S8C, D). (This is likely the reason why geneTrajectory under-performed in our test cases.) As we demonstrated, multiple gene sets contribute to the variation in the data, each providing a distance measure that differs from that of the other sets. A single cell– cell graph (such as that constructed by geneTrajectory and similar methods) cannot fully articulate complex, multi-faceted notions of cell–cell similarity.

In addition to identifying features that encode different topological structures, we found that ID can also be a powerful tool for integrative analysis. By applying ID on two independently collected scRNA-seq data of the human lung, we observed in each a set of genes that is sensitive to experimental covariates, a set of genes that defines a “barrel”-shaped differentiation trajectory of lung epithelial cells, and a final set of genes that charts cell cycle progression. We further showed that the differentiation and cell cycle related genes are highly conserved across donor and other experimental covariates. In the future, it will be worthwhile to employ the same strategy to investigate if there are other topological structures conserved across related datasets, and if the same structure is defined by the same set of genes.

ID constructs an alternative parametrization of the system by a variational autoencoder (VAE) [40] and identifies related features by looking for those that respond similarly to a small perturbation in the latent space. We believe ID is not the only way to identify these different gene sets. For example, simpler approaches can be used to achieve the same goal. In our analysis, we showed that UMAP of the transpose of the data matrix is sufficient when the similarity of cells defined by the different gene subsets differs dramatically (Figure 2). However, in more complex scenarios as shown in Figure 3, UMAP of the transpose of the data matrix becomes unable to separate features that encode distinct structures. We also showed that NMF performed well when the data is broken down into an optimal number of components. However, NMF becomes computationally costly as the number of components increases (Figure S4), and finding the optimal number of component is itself inefficient.

A critical assumption we have made in our analysis is that each gene only participates in a single structure. While this assumption should hold under most circumstances, it would be exciting to adapt ID for more complex datasets, such as those that contain both “normal” and “diseased” cells. In such cases, by analyzing “normal” and “diseased” cells separately, one can identify genes with context specific structures. For example, genes that used to be part of a “circle” may become part of a “branch” in diseased cells due to dysregulated gene expression.

In this work, we raised the issue that the relationship between cells is more complicated than their pairwise distances on the gene expression space, addressed it using ID, and demonstrated its broad utility across diverse datasets and biological systems. We envision that ID can be useful in a variety of ways. First, we demonstrate through multiple examples that ID–derived feature selection can support more informative data visualization. Second, by identifying genes associated with each structure, ID allows one to reduce the daunting task of characterizing the dynamics of a high-dimensional system to a much simpler problem of characterizing the dynamics of a small number of weakly coupled low-dimensional systems. This suggests that, unlike the existing paradigm of parametrizing dynamics on a single low-dimensional space [57, 58, 59], the correct approach to model the dynamics of a cell is to simultaneously evolve along multiple subspaces. This same strategy can also be adopted to characterize multi-modal data. Finally, it can be used to discriminate between technical artifacts and real biological signals, enabling batch correction. Altogether, ID provides a computational framework that can provide insights into dynamic gene regulation, predict how the system will respond to perturbations, and eventually guide the design of control strategies.

## 4 Methods

### 4.1 VAE optimization

By default, ID uses a 2–layer encoder neural network with 300 fully connected neurons per layer and ReLU activation to project count data onto a 50–dimensional latent space (**Z**). The data represented in **Z** are used as input to a decoder network to reconstruct the observed data 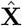. The loss function is simply the mean squared error between the input data **X** and the reconstructed data 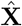. During each training loop, we focus only on a randomly selected small subset of the data, by default 150. In all test cases, the optimization of the model parameters was done with the ADAM [60] optimizer as implemented in pytorch [61] with a learning rate of 0.0005, *β*_1_ = 0.8, *β*_2_ = 0.9, and a weight decay of 0.0001. No scheduler was used to change the learning rate during the training process.

### 4.2 Application of alternative approaches

We compared ID to alternative approaches that can be used to decouple multiple components of variation in the data.

#### 4.2.1 Non-negative matrix factorization

Non-negative matrix factorization aims to deconstruct the original data matrix **X** into multiple components:

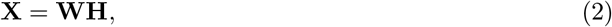

where 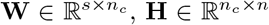 and *n*_*c*_ denotes the number of components the data matrix has been broken into. We used the matrix **H** to cluster genes in two ways. First, we used **H** directly, where each gene is defined by a vector of dimension *n*_*c*_. Alternatively, we first reduce the dimensionality of **H** before conducting clustering on the low-dimensional space. In all cases, NMF is implemented by `sklearn.decomposition.NMF` with default parameters.

#### 4.2.2 GeneTrajectory

GeneTrajectory [30] was applied with code downloaded from https://github.com/KlugerLab/GeneTrajectorywith default parameters.

#### 4.2.3 Constructing an embedding for genes using UMAP

While UMAP is typically used to visualize clusters of cells, it may also be used on the transpose of the data to visualize clusters of genes, which are potentially those that define similar structures in the data. We constructed an embedding for genes with UMAP by first transposing the data matrix, **X**, after normalization and extracting top 20 principal components before applying UMAP (**n_neighbour**=5).

#### 4.2.4 Applying clustering algorithms

K-means clustering and spectral clustering are applied using the scikit-learn [62] package in python with default parameters. A radial basis function kernel was used to construct the affinity matrix during spectral clustering using the top ten neighbors of each cell. Since the number of distinct structures is known for the toy datasets, we set the `n_clusters` parameter for both `sklearn.cluster.SpectralClustering` and `sklearn.cluster.Kmeans` to the number of distinct structures.

For real datasets where the number of clusters is not known, we applied DBSCAN using the scikitlearn [62] package to automatically conduct cluster assignment with `eps=0.5` and `min_samples=3`.

### 4.3 Datasets

We illustrate ID using a synthetic data, and then demonstrate it in a set of five real scRNA-seq datasets.

#### 4.3.1 Synthetic “toy” data

The toy datasets we used are constructed in two steps. First, we define the topology of the data on a low-dimensional space, which is taken to be the underlying manifold for the data. Then, via a topology preserving transformation, we lift the low-dimensional data into higher dimensions to mimic how the underlying low–dimensional structure is manifested in the high–dimensional gene expression space. For the “linear”, “2-branch”, and “4-branch” toy datasets, we define low-dimensional topology using the scattered dots illustrated in Figure S1B. Next, we use UMAP to project them from this low-dimensional space to a 50-dimensional space. For the toy data with complex topologies, we first sampled from the 3D shapes directly before projecting them into higher dimensions using UMAP. Raw data containing the mesh representation of these 3D shapes can be found at https://www.craftsmanspace.com/free-3d-models/topological-mesh-modeling-examples.html.

#### 4.3.2 Blood differentiation

The blood differentiation dataset [42, 41] was downloaded from https://github.com/HaghverdiLab/velocity_notebooks/tree/main/datasets/HSPC. This dataset contains 2430 cells with a mean UMI 17925. It has been enriched for long-term hematopoetic stem cells. Cells within this dataset have not reached final states of mature blood cells.

For this and all other real scRNA-seq datasets, we preprocessed the data by first normalizing each cell by its library size and then *Z*-scoring each feature. 3000 highly variable genes were selected by scanpy [63] prior to applying ID.

#### 4.3.3 Mouse hippocampus

The mouse hippocampus dataset [43, 44] was downloaded from http://pklab.med.harvard.edu/velocyto/DentateGyrus. It contains 18213 cells with a mean UMI of 4677. Preprocessing was performed as above.

#### 4.3.4 Mouse microglia

The mouse microgila dataset [45] was downloaded from https://cellxgene.cziscience.com/collections/4828d33d-fb26-42e7-bf36-18293b0eec85. This dataset contained 12330 sorted microglia with a mean UMI of 3682. Preprocessing was performed as above.

#### 4.3.5 Mouse hair follicle

The mouse hair follicle dataset [30] was downloaded from https://figshare.com/articles/dataset/Processed_Seurat_objects_for_GeneTrajectory_inference_Gene_Trajectory_Inference_for_Single-cell_Data_by_Optimal_Transport_Metrics_/25243225. The dataset contains both wild type cells and cells with Wnt knock-out. It contains a total of 10328 cells with a mean UMI of 13948. Preprocessing was performed as above.

#### 4.3.6 Human lung samples

The human lung organoid data [55] was downloaded from https://cellxgene.cziscience.com/collections/e9cf4e8d-05ed-4d95-b550-af35ca219390. It is collected from two separate human donors, dissociated and used to generate lung organoid (O1). O1 derived transitionally differentiated (TD) cells are cultured in lung organoid growth medium, PneumaCult EX-plus medium and PneumaCult-ALI maintenance medium to generate the O2, EX and the ALIEX model. This data contains 15806 cells with a mean UMI of 3833. Preprocessing was performed as above.

The human lung sample data [54] was downloaded from https://cellxgene.cziscience.com/collections/73cf6939-3caa-4105-bc57-e073ee885a28. It is obtained from two human transplant donor lungs that are considered unsuitable for transplantation. Cells are isolated from the bronchial rings, dissociated and then cultured in four different culture media. This dataset contains 10224 cells with a mean UMI 2826. Preprocessing was performed as above.

#### 4.3.7 Human HSCs

The human HSC dataset, sampled from three human donors, was downloaded from [56]. The dataset contains log-normalized counts only. For this dataset, we first selected the top 2000 genes and rescaled log-normalized count data via Z-scoring.

### 4.4 Quantifying the extent of overlap between gene sets

Suppose *G*_1_ and *G*_2_ are the features (total genes) of dataset 1 and dataset 2 respectively. We are interested in quantifying whether overlap between an ID-derived subset of *G*_1_, denoted as *S*_1_, and an ID-derived subset of *G*_2_, *S*_2_, is greater than expected by chance. In the case where *G*_1_ = *G*_2_, the probability distribution for the overlap |*S*_1_ ∩ *S*_2_| is hypergeometric. However, when the base feature sets *G*_1_ and *G*_2_ have differences, the distribution becomes more complicated (see Supplementary Text). To construct this distribution, we randomly sample |*S*_1_| genes from *G*_1_ and |*S*_2_| genes from *G*_2_ to obtain two new gene sets 𝒮′_1_ and 𝒮′_2_. We repeat this 15,000 times to obtain a distribution of |𝒮′_1_ ∩ 𝒮′_2_|, i.e. the expected size of the overlap between |*S*_1_| and |*S*_2_| if they had been drawn at random from *G*_1_ and *G*_2_. A *p*-value for the observed |*S*_1_ ∩ *S*_2_| may then be obtained from this distribution. We note that an exact *p*-value may also be calculated, as given in the Supplementary Text; however, these calculations must be undertaken with care to be efficient and stable, and so we opted to bootstrap it instead.

## 5 Acknowledgements

This work was supported by NSF grant DMS-1764421, Simons Foundation grant 597491, and NIH grant R01AG068579.

## 6 Code availability

The code used for this manuscript can be found in https://github.com/bxxu/Identification-of-distinct-topology

## References

[1] C.H. Waddington. The Strategy of the Genes. Routledge, 1957.

[2] Caleb Weinreb, Alejo Rodriguez-Fraticelli, Fernando D. Camargo, and Allon M. Klein. Lineage tracing on transcriptional landscapes links state to fate during differentiation. Science, 367(6479):eaaw3381, February 2020.

[3] Simon L. Freedman, Bingxian Xu, Sidhartha Goyal, and Madhav Mani. A dynamical systems treatment of transcriptomic trajectories in hematopoiesis. Development, 150(11):dev201280, June 2023.

[4] J. M. W. Slack. From Egg to Embryo: Regional Specification in Early Development. Cambridge University Press, 2 edition, May 1991.

[5] Zhilin Qu, W. Robb MacLellan, and James N. Weiss. Dynamics of the Cell Cycle: Checkpoints, Sizers, and Timers. Biophysical Journal, 85(6):3600–3611, December 2003.

[6] Timothy S. Gardner, Milos Dolnik, and James J. Collins. A theory for controlling cell cycle dynamics using a reversibly binding inhibitor. Proceedings of the National Academy of Sciences, 95(24):14190–14195, November 1998.

[7] Andrea Riba, Attila Oravecz, Matej Durik, Sara Jiménez, Violaine Alunni, Marie Cerciat, Matthieu Jung, Céline Keime, William M. Keyes, and Nacho Molina. Cell cycle gene regulation dynamics revealed by RNA velocity and deep-learning. Nature Communications, 13(1):2865, May 2022.

[8] Wayne Stallaert, Katarzyna M. Kedziora, Colin D. Taylor, Tarek M. Zikry, Jolene S. Ranek, Holly K. Sobon, Sovanny R. Taylor, Catherine L. Young, Jeanette G. Cook, and Jeremy E. Purvis. The structure of the human cell cycle. Cell Systems, 13(3):230–240.e3, March 2022.

[9] Ravi Allada and Brian Y. Chung. Circadian Organization of Behavior and Physiology in Drosophila. Annual Review of Physiology, 72(1):605–624, March 2010.

[10] Ray Zhang, Nicholas F. Lahens, Heather I. Ballance, Michael E. Hughes, and John B. Hogenesch. A circadian gene expression atlas in mammals: Implications for biology and medicine. Proceedings of the National Academy of Sciences, 111(45):16219–16224, November 2014.

[11] John S. O’Neill and Akhilesh B. Reddy. Circadian clocks in human red blood cells. Nature, 469(7331):498–503, January 2011.

[12] Bingxian Xu, Dae-Sung Hwangbo, Sumit Saurabh, Clark Rosensweig, Ravi Allada, William L. Kath, and Rosemary Braun. Temperature-driven coordination of circadian transcriptional regulation. PLOS Computational Biology, 20(4):e1012029, April 2024.

[13] Chunghun Lim and Ravi Allada. Emerging roles for post-transcriptional regulation in circadian clocks. Nature Neuroscience, 16(11):1544–1550, November 2013.

[14] Leland McInnes, John Healy, and James Melville. UMAP: Uniform Manifold Approximation and Projection for Dimension Reduction, September 2020. arXiv:1802.03426 [stat].

[15] Laleh Haghverdi, Maren Büttner, F Alexander Wolf, Florian Buettner, and Fabian J Theis. Diffusion pseudotime robustly reconstructs lineage branching. Nature Methods, 13(10):845–848, October 2016.

[16] Laurens van der Maaten and Geoffrey Hinton. Visualizing data using t-sne. Journal of Machine Learning Research, 9(86):2579–2605, 2008.

[17] Xiaojie Qiu, Qi Mao, Ying Tang, Li Wang, Raghav Chawla, Hannah A Pliner, and Cole Trapnell. Reversed graph embedding resolves complex single-cell trajectories. Nature Methods, 14(10):979– 982, October 2017.

[18] Jiarui Ding and Aviv Regev. Deep generative model embedding of single-cell RNA-Seq profiles on hyperspheres and hyperbolic spaces. Nature Communications, 12(1):2554, May 2021.

[19] Jiarui Ding, Anne Condon, and Sohrab P. Shah. Interpretable dimensionality reduction of single cell transcriptome data with deep generative models. Nature Communications, 9(1):2002, May 2018.

[20] Daniel Lee and H. Sebastian Seung. Algorithms for Non-negative Matrix Factorization. In T. Leen, T. Dietterich, and V. Tresp, editors, Advances in Neural Information Processing Systems, volume 13. MIT Press, 2000.

[21] Daniel D. Lee and H. Sebastian Seung. Learning the parts of objects by non-negative matrix factorization. Nature, 401(6755):788–791, October 1999.

[22] Pentti Paatero, Unto Tapper, Pasi Aalto, and Markku Kulmala. Matrix factorization methods for analysing diffusion battery data. Journal of Aerosol Science, 22:S273–S276, 1991.

[23] Pentti Paatero and Unto Tapper. Positive matrix factorization: A non-negative factor model with optimal utilization of error estimates of data values. Environmetrics, 5(2):111–126, June 1994.

[24] P Anttila, P Paatero, U Tapper, and O Jarvinen. Source identification of bulk wet deposition in Finland by positive matrix factorization. Atmospheric Environment, 29(14):1705–1718, 1995.

[25] Hanna Mendes Levitin, Jinzhou Yuan, Yim Ling Cheng, Francisco Jr Ruiz, Erin C Bush, Jeffrey N Bruce, Peter Canoll, Antonio Iavarone, Anna Lasorella, David M Blei, and Peter A Sims. De novo gene signature identification from single-cell RNA-seq with hierarchical Poisson factorization. Molecular Systems Biology, 15(2):e8557, February 2019.

[26] Karin Pelka, Matan Hofree, Jonathan H. Chen, Siranush Sarkizova, Joshua D. Pirl, Vjola Jorgji, Alborz Bejnood, Danielle Dionne, William H. Ge, Katherine H. Xu, Sherry X. Chao, Daniel R. Zollinger, David J. Lieb, Jason W. Reeves, Christopher A. Fuhrman, Margaret L. Hoang, Toni Delorey, Lan T. Nguyen, Julia Waldman, Max Klapholz, Isaac Wakiro, Ofir Cohen, Julian Albers, Christopher S. Smillie, Michael S. Cuoco, Jingyi Wu, Mei-ju Su, Jason Yeung, Brinda Vijaykumar, Angela M. Magnuson, Natasha Asinovski, Tabea Moll, Max N. Goder-Reiser, Anise S. Applebaum, Lauren K. Brais, Laura K. DelloStritto, Sarah L. Denning, Susannah T. Phillips, Emma K. Hill, Julia K. Meehan, Dennie T. Frederick, Tatyana Sharova, Abhay Kanodia, Ellen Z. Todres, Judit Jané-Valbuena, Moshe Biton, Benjamin Izar, Conner D. Lambden, Thomas E. Clancy, Ronald Bleday, Nelya Melnitchouk, Jennifer Irani, Hiroko Kunitake, David L. Berger, Amitabh Srivastava, Jason L. Hornick, Shuji Ogino, Asaf Rotem, Sébastien Vigneau, Bruce E. Johnson, Ryan B. Corcoran, Arlene H. Sharpe, Vijay K. Kuchroo, Kimmie Ng, Marios Giannakis, Linda T. Nieman, Genevieve M. Boland, Andrew J. Aguirre, Ana C. Anderson, Orit Rozenblatt-Rosen, Aviv Regev, and Nir Hacohen. Spatially organized multicellular immune hubs in human colorectal cancer. Cell, 184(18):4734–4752.e20, September 2021.

[27] Russell Z. Kunes, Thomas Walle, Max Land, Tal Nawy, and Dana Pe’er. Supervised discovery of interpretable gene programs from single-cell data. Nature Biotechnology, 42(7):1084–1095, July 2024.

[28] Sarah Chen, Aviv Regev, Anne Condon, and Jiarui Ding. CellUntangler: separating distinct biological signals in single-cell data with deep generative models, January 2025.

[29] Jonathan Karin, Yonathan Bornfeld, and Mor Nitzan. scPrisma infers, filters and enhances topological signals in single-cell data using spectral template matching. Nature Biotechnology, 41(11):1645–1654, November 2023.

[30] Rihao Qu, Xiuyuan Cheng, Esen Sefik, Jay S. Stanley Iii, Boris Landa, Francesco Strino, Sarah Platt, James Garritano, Ian D. Odell, Ronald Coifman, Richard A. Flavell, Peggy Myung, and Yuval Kluger. Gene trajectory inference for single-cell data by optimal transport metrics. Nature Biotechnology, 43(2):258–268, February 2025.

[31] Cole Trapnell, Davide Cacchiarelli, Jonna Grimsby, Prapti Pokharel, Shuqiang Li, Michael Morse, Niall J Lennon, Kenneth J Livak, Tarjei S Mikkelsen, and John L Rinn. The dynamics and regulators of cell fate decisions are revealed by pseudotemporal ordering of single cells. Nature Biotechnology, 32(4):381–386, April 2014.

[32] Junyue Cao, Malte Spielmann, Xiaojie Qiu, Xingfan Huang, Daniel M. Ibrahim, Andrew J. Hill, Fan Zhang, Stefan Mundlos, Lena Christiansen, Frank J. Steemers, Cole Trapnell, and Jay Shendure. The single-cell transcriptional landscape of mammalian organogenesis. Nature, 566(7745):496–502, February 2019.

[33] Jiawei Huang, Jie Sheng, and Daifeng Wang. Manifold learning analysis suggests strategies to align single-cell multimodal data of neuronal electrophysiology and transcriptomics. Communications Biology, 4(1):1308, November 2021.

[34] Charles Fefferman, Sanjoy Mitter, and Hariharan Narayanan. Testing the manifold hypothesis. Journal of the American Mathematical Society, 29(4):983–1049, February 2016.

[35] Juan A. Gallego, Matthew G. Perich, Lee E. Miller, and Sara A. Solla. Neural Manifolds for the Control of Movement. Neuron, 94(5):978–984, June 2017.

[36] Vasyl Alba, James E Carthew, Richard W Carthew, and Madhav Mani. Global constraints within the developmental program of the Drosophila wing. eLife, 10:e66750, June 2021.

[37] Giulio Bondanelli, Thomas Deneux, Brice Bathellier, and Srdjan Ostojic. Network dynamics underlying OFF responses in the auditory cortex. eLife, 10:e53151, March 2021.

[38] Hansem Sohn, Devika Narain, Nicolas Meirhaeghe, and Mehrdad Jazayeri. Bayesian Computation through Cortical Latent Dynamics. Neuron, 103(5):934–947.e5, September 2019.

[39] Peiran Gao, Eric Trautmann, Byron Yu, Gopal Santhanam, Stephen Ryu, Krishna Shenoy, and Surya Ganguli. A theory of multineuronal dimensionality, dynamics and measurement, November 2017.

[40] Diederik P Kingma and Max Welling. Auto-Encoding Variational Bayes, 2013. Version Number: 11.

[41] Yasmin Demerdash, Brigitte Bouman, Laleh Haghverdi, and Marieke Essers. Unbiased, longitudinal analysis of the inflammatory response of hspcs at the single cell level resolves controversies regarding the HSPCs stress response. Experimental Hematology, 111:S80, 2022.

[42] Valérie Marot-Lassauzaie, Brigitte Joanne Bouman, Fearghal Declan Donaghy, Yasmin Demerdash, Marieke Alida Gertruda Essers, and Laleh Haghverdi. Towards reliable quantification of cell state velocities. PLOS Computational Biology, 18(9):e1010031, September 2022.

[43] Gioele La Manno, Ruslan Soldatov, Amit Zeisel, Emelie Braun, Hannah Hochgerner, Viktor Petukhov, Katja Lidschreiber, Maria E. Kastriti, Peter Lönnerberg, Alessandro Furlan, Jean Fan, Lars E. Borm, Zehua Liu, David Van Bruggen, Jimin Guo, Xiaoling He, Roger Barker, Erik Sundström, Gonçalo Castelo-Branco, Patrick Cramer, Igor Adameyko, Sten Linnarsson, and Peter V. Kharchenko. RNA velocity of single cells. Nature, 560(7719):494–498, August 2018.

[44] Hannah Hochgerner, Amit Zeisel, Peter Lönnerberg, and Sten Linnarsson. Conserved properties of dentate gyrus neurogenesis across postnatal development revealed by single-cell RNA sequencing. Nature Neuroscience, 21(2):290–299, February 2018.

[45] Caroline C. Escoubas, Leah C. Dorman, Phi T. Nguyen, Christian Lagares-Linares, Haruna Nakajo, Sarah R. Anderson, Jerika J. Barron, Sarah D. Wade, Beatriz Cuevas, Ilia D. Vainchtein, Nicholas J. Silva, Ricardo Guajardo, Yinghong Xiao, Peter V. Lidsky, Ellen Y. Wang, Brianna M. Rivera, Sunrae E. Taloma, Dong Kyu Kim, Elizaveta Kaminskaya, Hiromi Nakao-Inoue, Bjoern Schwer, Thomas D. Arnold, Ari B. Molofsky, Carlo Condello, Raul Andino, Tomasz J. Nowakowski, and Anna V. Molofsky. Type-I-interferon-responsive microglia shape cortical development and behavior. Cell, 187(8):1936–1954.e24, April 2024.

[46] Ainhoa Plaza-Zabala, Virginia Sierra-Torre, and Amanda Sierra. Autophagy and Microglia: Novel Partners in Neurodegeneration and Aging. International Journal of Molecular Sciences, 18(3):598, March 2017.

[47] Fushan Shi, Yang Yang, Mohammed Kouadir, Yongyao Fu, Lifeng Yang, Xiangmei Zhou, Xiaomin Yin, and Deming Zhao. Inhibition of phagocytosis and lysosomal acidification suppresses neurotoxic prion peptide-induced NALP3 inflammasome activation in BV2 microglia. Journal of Neuroimmunology, 260(1–2):121–125, July 2013.

[48] Harini Iyer, Kimberle Shen, Ana M. Meireles, and William S. Talbot. A lysosomal regulatory circuit essential for the development and function of microglia. Science Advances, 8(35):eabp8321, September 2022.

[49] Peggy Myung, Thomas Andl, and Radhika Atit. The origins of skin diversity: lessons from dermal fibroblasts. Development, 149(23):dev200298, December 2022.

[50] Rihao Qu, Khusali Gupta, Danni Dong, Yiqun Jiang, Boris Landa, Charles Saez, Gwendolyn Strickland, Jonathan Levinsohn, Pei-lun Weng, M. Mark Taketo, Yuval Kluger, and Peggy Myung. Decomposing a deterministic path to mesenchymal niche formation by two intersecting morphogen gradients. Developmental Cell, 57(8):1053–1067.e5, April 2022.

[51] Khusali Gupta, Jonathan Levinsohn, George Linderman, Demeng Chen, Thomas Yang Sun, Danni Dong, M. Mark Taketo, Marcus Bosenberg, Yuval Kluger, Keith Choate, and Peggy Myung. Single-Cell Analysis Reveals a Hair Follicle Dermal Niche Molecular Differentiation Trajectory that Begins Prior to Morphogenesis. Developmental Cell, 48(1):17–31.e6, January 2019.

[52] Jiang Fu and Wei Hsu. Epidermal Wnt Controls Hair Follicle Induction by Orchestrating Dynamic Signaling Crosstalk between the Epidermis and Dermis. Journal of Investigative Dermatology, 133(4):890–898, April 2013.

[53] Beate M. Lichtenberger, Maria Mastrogiannaki, and Fiona M. Watt. Epidermal β-catenin activation remodels the dermis via paracrine signalling to distinct fibroblast lineages. Nature Communications, 7(1):10537, February 2016.

[54] Elisa Redman, Morgane Fierville, Amélie Cavard, Magali Plaisant, Marie-Jeanne Arguel, Sandra Ruiz Garcia, Eamon M. McAndrew, Cédric Girard-Riboulleau, Kevin Lebrigand, Virginie Magnone, Gilles Ponzio, Delphine Gras, Pascal Chanez, Sophie Abelanet, Pascal Barbry, Brice Marcet, and Laure-Emmanuelle Zaragosi. Cell Culture Differentiation and Proliferation Conditions Influence the In Vitro Regeneration of the Human Airway Epithelium. American Journal of Respiratory Cell and Molecular Biology, 71(3):267–281, September 2024.

[55] Jieun Kim, Eun-Young Eo, Bokyong Kim, Heetak Lee, Jihoon Kim, Bon-Kyoung Koo, Hyung-Jun Kim, Sukki Cho, Jinho Kim, and Young-Jae Cho. Transcriptomic Analysis of Air–Liquid Interface Culture in Human Lung Organoids Reveals Regulators of Epithelial Differentiation. Cells, 13(23):1991, December 2024.

[56] Daniel Burkhardt, Malte Luecken, Andrew Benz, Peter Holderrieth, Jonathan Bloom, Christopher Lance, Ashley Chow, and Ryan Holbrook. Open problems - multimodal single-cell integration. https://kaggle.com/competitions/open-problems-multimodal, 2022. Kaggle.

[57] Bingxian Xu and Rosemary Braun. VIST: variational inference for single cell time series. Genome Biology, 27(1):29, January 2026.

[58] Alexander Tong, Jessie Huang, Guy Wolf, David van Dijk, and Smita Krishnaswamy. TrajectoryNet: A Dynamic Optimal Transport Network for Modeling Cellular Dynamics. arχiv preprint, 2020. Publisher: arXiv Version Number: 2.

[59] Grace Hui Ting Yeo, Sachit D. Saksena, and David K. Gifford. Generative modeling of single-cell time series with PRESCIENT enables prediction of cell trajectories with interventions. Nature Communications, 12(1):3222, May 2021.

[60] Diederik P. Kingma and Jimmy Ba. Adam: A Method for Stochastic Optimization, January 2017. arXiv:1412.6980 [cs].

[61] Adam Paszke, Sam Gross, Francisco Massa, Adam Lerer, James Bradbury, Gregory Chanan, Trevor Killeen, Zeming Lin, Natalia Gimelshein, Luca Antiga, Alban Desmaison, Andreas Köpf, Edward Yang, Zach DeVito, Martin Raison, Alykhan Tejani, Sasank Chilamkurthy, Benoit Steiner, Lu Fang, Junjie Bai, and Soumith Chintala. PyTorch: An Imperative Style, High-Performance Deep Learning Library, December 2019. arXiv:1912.01703 [cs, stat].

[62] Fabian Pedregosa, Gaël Varoquaux, Alexandre Gramfort, Vincent Michel, Bertrand Thirion, Olivier Grisel, Mathieu Blondel, Peter Prettenhofer, Ron Weiss, Vincent Dubourg, Jake Vanderplas, Alexandre Passos, David Cournapeau, Matthieu Brucher, Matthieu Perrot, and Édouard Duchesnay. Scikit-learn: Machine learning in python. J. Mach. Learn. Res., 12(null):2825–2830, November 2011.

[63] F. Alexander Wolf, Philipp Angerer, and Fabian J. Theis. SCANPY: large-scale single-cell gene expression data analysis. Genome Biology, 19(1):15, December 2018.

