## supplementary figures for "Identification of Distinct Topological Structures From High-Dimensional Data"

### 7 Supplemental material

#### S1 Exact quantification of the extent of overlap between gene sets

Suppose $G_1$ and $G_2$ are the features of dataset 1 and dataset 2 respectively, having sizes $N_1 = |G_1|$ and $N_2 = |G_2|$. Notably, $G_1$ and $G_2$ are not identical, although they do overlap; we denote the size of the overlap as $N^* = |G_1 \cap G_2|$.

We are interested in quantifying the extent to which ID-derived subsets $S_1$ (from $G_1$) and $S_2$ (from $G_2$) overlap. Denote the sizes of $S_1$ and $S_2$ as $m_1 = |S_1|$ and $m_2 = |S_2|$, and the size of their intersection as $m^* = |S_1 \cap S_2|$. Essentially, we would like to know the probability of the overlap $m^*$, $\Pr(m^*; N_1, N_2, N^*, m_1, m_2)$, between two draws of size $m_1$ and $m_2$ from populations of size $N_1$ and $N_2$ that have an overlap of size $N^*$.

It is clear that because $G_1 \neq G_2$, this is not a simple hypergeometric distribution; rather, we seeking the distribution of the intersection size of two independent hypergeometric samples drawn from two *partially* overlapping finite populations. Any draws from $G_1$ or $G_2$ that do not hit the overlap set $G_1 \cap G_2$ have no chance of hitting each other. We can thus decompose the probability into the probability that the $m_1$ draws hit the overlap set of size $N^*$, the probability that the $m_2$ draws hit $N^*$, and the probability that the draws within the overlap set hit each other, conditioned on the number of draws in that hit the overlap set. Let us denote $k_1$ as the number of draws that hit the overlap, i.e. $k_1 = |S_1 \cap (G_1 \cap G_2)|$ and likewise $k_2 = |S_2 \cap (G_1 \cap G_2)|$. We can decompose the probability as

$$\Pr(m^*; N_1, N_2, N^*, m_1, m_2) = \\ = \sum_{k_1=0}^{\min(m_1,N^*)} \sum_{k_2=0}^{\min(m_2,N^*)} \underbrace{\Pr(k_1; N_1, N^*, m_1)}_{\substack{\text{probability of } S_1 \\ \text{hitting overlap } G_1 \cap G_2}} \underbrace{\Pr(k_2; N_2, N^*, m_2)}_{\substack{\text{probability of } S_2 \\ \text{hitting overlap } G_1 \cap G_2}} \underbrace{\Pr(m^*; N^*, k_1, k_2)}_{\substack{\text{probability of the overlap} \\ \text{``hits'' intersecting}}} . \tag{S1}$$

For the last term to be nonzero, $k_1$ and $k_2$ must be at least $m^*$ (it is impossible to have an intersection that exceeds the size the draw, and thus we can write the sums starting from $m^*$. In what follows, we will further assume that $m_1, m_2 < N^*$, such that the sizes $k_1, k_2$ are limited by $m_1, m_2$ and not $N^*$. The RHS of Eq. S1 is simply the product of hypergeometric probabilities,

$$= \sum_{k_1=m^*}^{m_1} \sum_{k_2=m^*}^{m_2} \frac{\binom{N^*}{k_1}\binom{N_1 - N^*}{m_1 - k_1}}{\binom{N_1}{m_1}} \frac{\binom{N^*}{k_2}\binom{N_2 - N^*}{m_2 - k_2}}{\binom{N_2}{m_2}} \frac{\binom{k_1}{m^*}\binom{N^* - k_1}{k2 - m^*}}{\binom{N^*}{k2}} \tag{S2}$$

$$= \frac{1}{\binom{N_1}{m_1}\binom{N_2}{m_2}} \sum_{k_1=m^*}^{m_1} \binom{N^*}{k_1}\binom{N_1 - N^*}{m_1 - k_1}\binom{k_1}{m^*}\left[\sum_{k_2=m^*}^{m_2} \binom{N_2 - N^*}{m_2 - k_2}\binom{N^* - k_1}{k_2 - m^*}\right] \tag{S3}$$

$$= \frac{1}{\binom{N_1}{m_1}\binom{N_2}{m_2}} \sum_{k_1=m^*}^{m_1} \binom{N^*}{k_1}\binom{N_1 - N^*}{m_1 - k_1}\binom{k_1}{m^*}\binom{N_2 - k_1}{m_2 - m^*} , \tag{S4}$$

where we have exploited the Vandermonde identity to reduce the sum over $k_2$. Nevertheless, the sum over $k_1$ remains. Rather than explicitly calculating the binomial coefficients for each term of the sum, one may compute it recursively as follows. Define

$$t_x = \binom{N^*}{x}\binom{N_1 - N^*}{m_1 - x}\binom{x}{m^*}\binom{N_2 - x}{m_2 - m^*}, \tag{S5}$$

where $x$ is used in place of $k_1$ above. Then, observing that $\binom{r}{x+1}/\binom{r}{x} = \frac{r-x}{x+1}$ and $\binom{x+1}{q}/\binom{x}{q} = \frac{x+1}{x-q+1}$ for any $r \geq (x+1)$ and $q \leq x$, we have

$$t_{x+1} = \frac{N^* - x}{x+1} \cdot \frac{m_1 - x}{(N_1 - N^*) - (m_1 - x) + 1} \cdot \frac{x+1}{x - m^* + 1} \cdot \frac{(N_2 - x) - (m_2 - m^*)}{N_2 - x} \cdot t_x\ , \tag{S6}$$

which can be further simplified slightly by canceling the $x+1$ terms. We begin the sum by computing the base case $x = m^*$

$$t_0 = t_{m^*} = \binom{N^*}{m^*}\binom{N_1 - N^*}{m_1 - m^*}\binom{N_2 - m^*}{m_2 - m^*} \tag{S7}$$

and then apply Eq. S6 to compute the successive terms up to $m_1$.

Together, Eqs. S4, S7 and S6 provide a means to calculate the PMF for the overlap of two samples $S_1$, $S_2$ from two partially–overlapping finite sets $G_1$, $G_2$; the cumulative probability, and hence a $p$-value, can be computed by summing the upper tail probabilities starting at $m^*$ and going to the maximal possible overlap, i.e. the minimum of $m_1$ and $m_2$ (again for simplicity assuming $m_1, m_2 < N^*$):

$$\begin{aligned}\Pr(M \geq m^*; N_1, N_2, N^*, m_1, m_2) &= \\ &= \frac{1}{\binom{N_1}{m_1}\binom{N_2}{m_2}} \sum_{m=m^*}^{\min(m_1,m_2)} \sum_{x=m}^{m_1} \binom{N^*}{x}\binom{N_1 - N^*}{m_1 - x}\binom{x}{m}\binom{N_2 - x}{m_2 - m}.\end{aligned} \tag{S8}$$

While this can be calculated exactly for moderate $N_1, N_2, N^*, m_1, m_2$, there is a risk of numerical instability due to integer overflow and catastrophic cancellation when $N_1, N_2, N^*$ as well as $m_1$ and $m_2$ are large. Two special–case simplifications are possible. First, in the case where $G_1 = G_2$ (and hence $N_1{=}N_2{=}N^*{=}N$), Eq. S1 reduces to a single hypergeometric distribution, $m^* \sim \text{Hypergeometric}(N, m_1, m_2)$. Second, in the case where $m_1$ and $m_2$ are small relative to $N_1$ and $N_2$ (sparse sampling), we can approximate Eq. S1 by considering

$$m^* = \sum_{i=1}^{N^*} \mathbb{1}_i\ , \tag{S9}$$

where the indicator function $\mathbb{1}_i = 1$ if shared label $i$ is drawn from both urns ($G_1$ and $G_2$). In the large $N_1, N_2$ limit, we can ignore the fact that we are sampling $m_1$ and $m_2$ without replacement and treat each term in the sum above (i.e., each draw) as independent draws from the same large underlying population. Thus, each label $i$ is selected with (approximate) probability $m_1/N_1$, $m_2/N_2$ for each urn, and the probability of a shared label is approximately $\Pr(\mathbb{1}_i{=}1) = \frac{m_1}{N_1}\frac{m_2}{N_2}$. In this regime, the shared labels behave as Bernoulli trials, of which we have $N^*$. Thus in the limit, Eq. S1 converges in distribution to

$$m^* \sim \text{Binomial}\left(N^*, \frac{m_1}{N_1}\frac{m_2}{N_2}\right). \tag{S10}$$

However, when $m_1$ and $m_2$ are large relative to $N_1$ and $N_2$, the independence assumption no longer applies and the approximation (Eq. S10) does not hold. Rather than computing the $p$-value exactly, we bootstrap it from the data as described in the Methods.

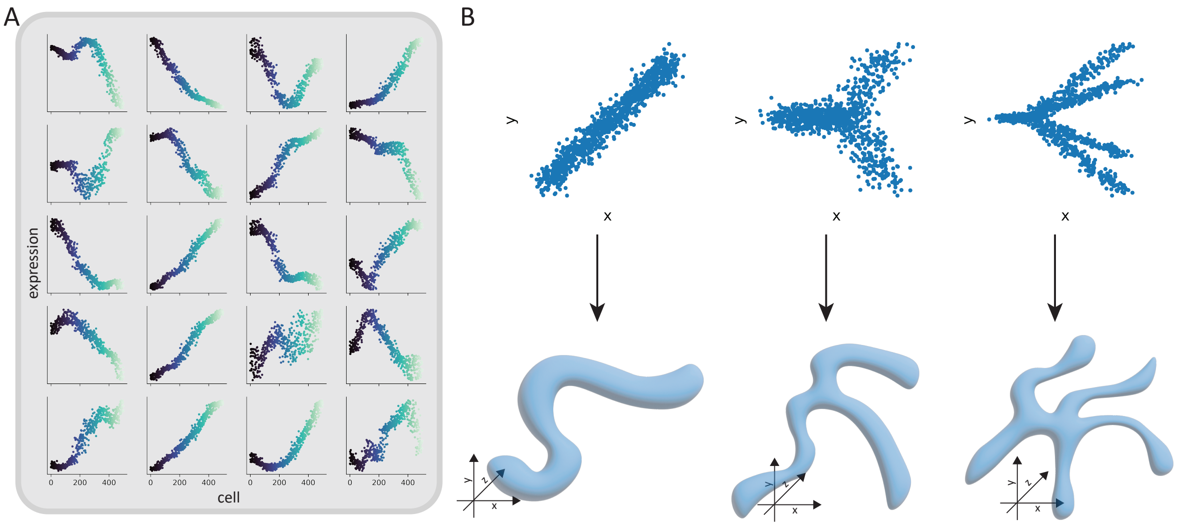


Figure S1: A: Example data constructed from an open curve (topological line). The initial topology is defined in 2D and projected to a 20-dimensional space. B: Illustration of the topology of the "linear", "branched" and the "4-branch" structures used Figure 2. These two-dimensional structures are then projected to a higher dimensional space.

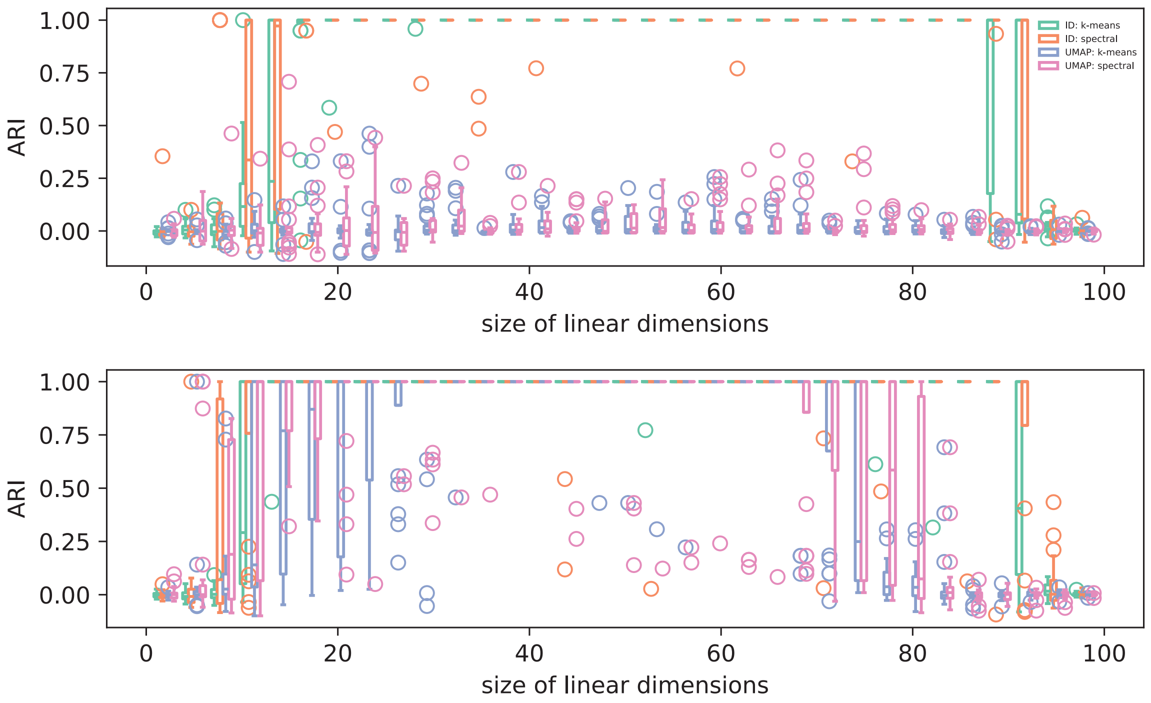


Figure S2: The full boxplot from Figure 2. In the top panel, the two processes had a similar order. In the bottom panel, the two processes had a vastly different order.

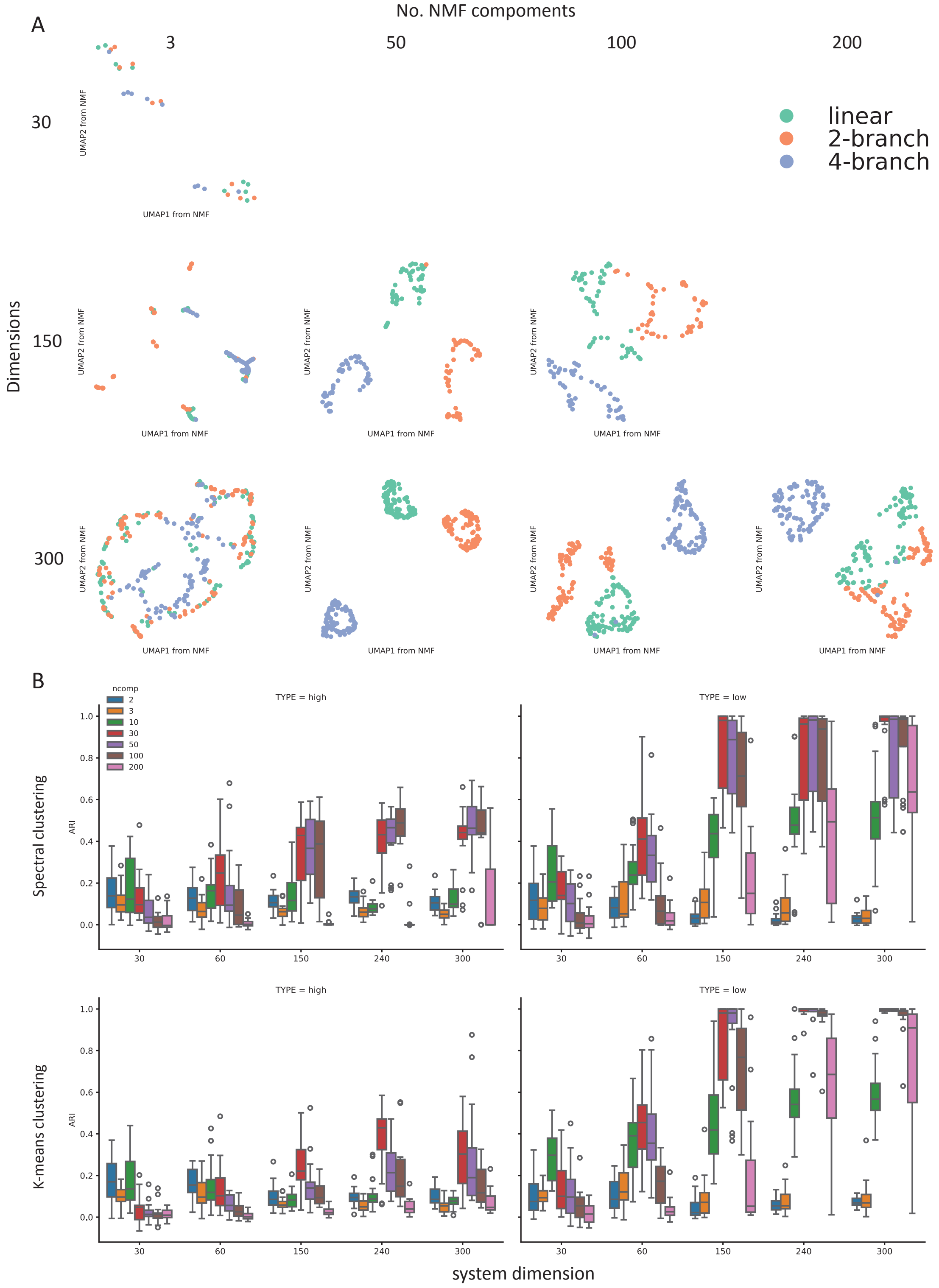


Figure S3: A: UMAP of genes constructed using results from NMF. The total dimension of the system used in the first, second and third row is 30, 150 and 300 respectively. The number of components the gene expression matrix is broken into in the first, second, third and forth column is 3, 50, 100 and 200 respectively. Plots were left blank when the number of NMF component exceeds the dimensionality of the system. B: ARI computed by clustering genes using spectral clustering (top panel) and $k$-means clustering (bottom) when the gene expression matrix is broken down to various numbers of components. **TYPE=high** indicates that clustering is done using all output from NMF, whereas **TYPE=low** indicates that NMF result is first projected to a 2D space using UMAP before clustering. Each trial was repeated 20 times.

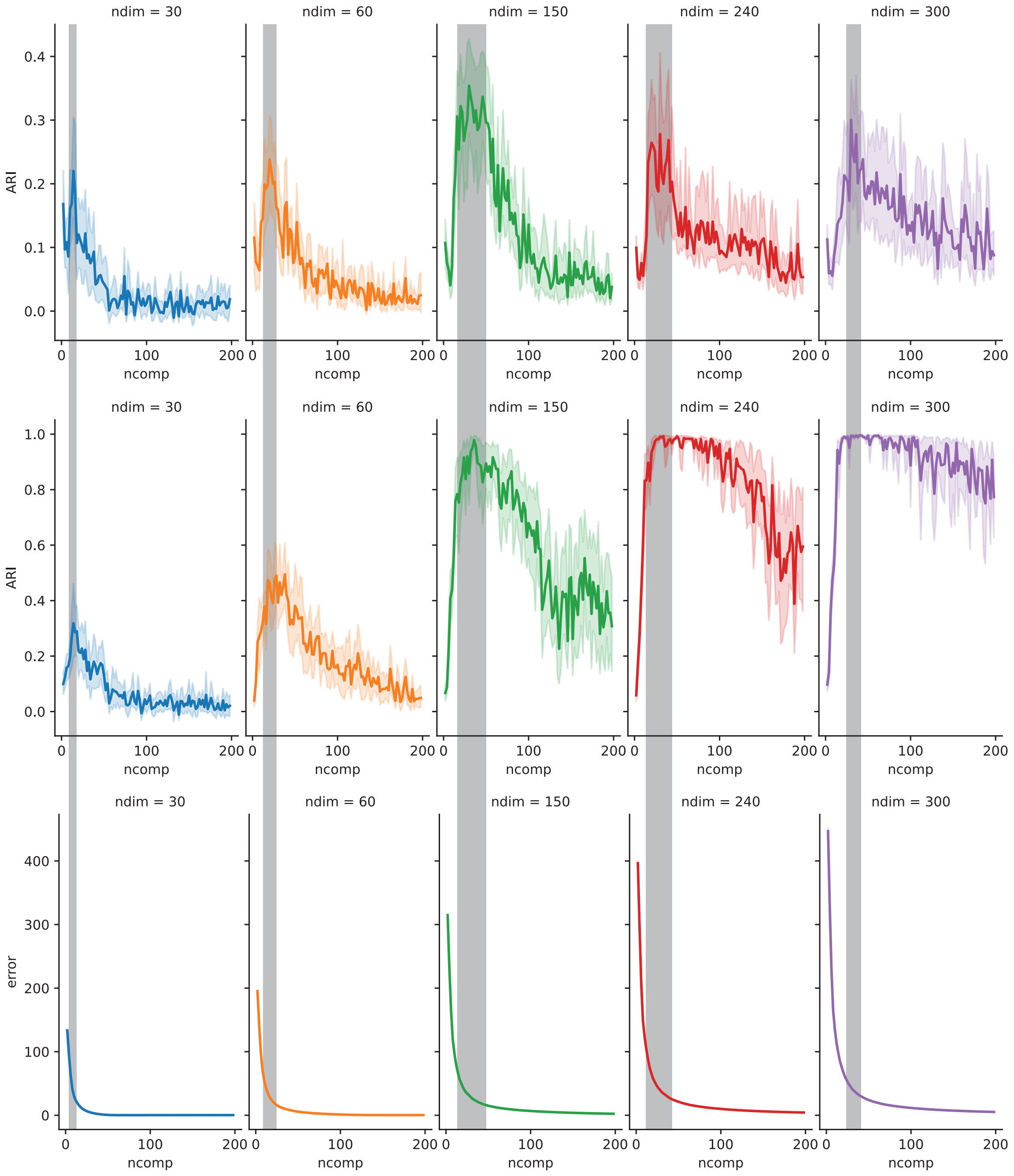


Figure S4: NMF is applied to the toy dataset from Figure 2D. ARIs were computed using $k$-means clustering in either the high-dimensional space (first row) or the low-dimensional space (second row) as both the system size (**ndim**) and the number of components (**ncomp**) varied. The third row illustrates how the NMF error changes with the number of components. The shading denotes the region where the ARI peak occurs when clustering is done in the high-dimensional space. Each trial was repeated 10 times.

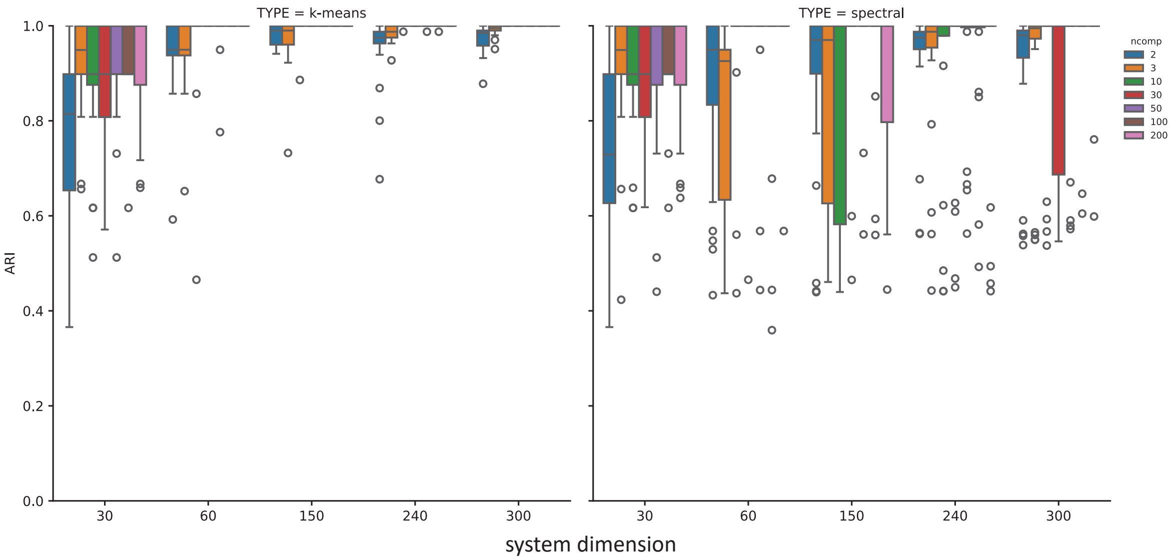


Figure S5: ARI computed using ID under various latent space dimensions using both $k$-means and spectral clustering.

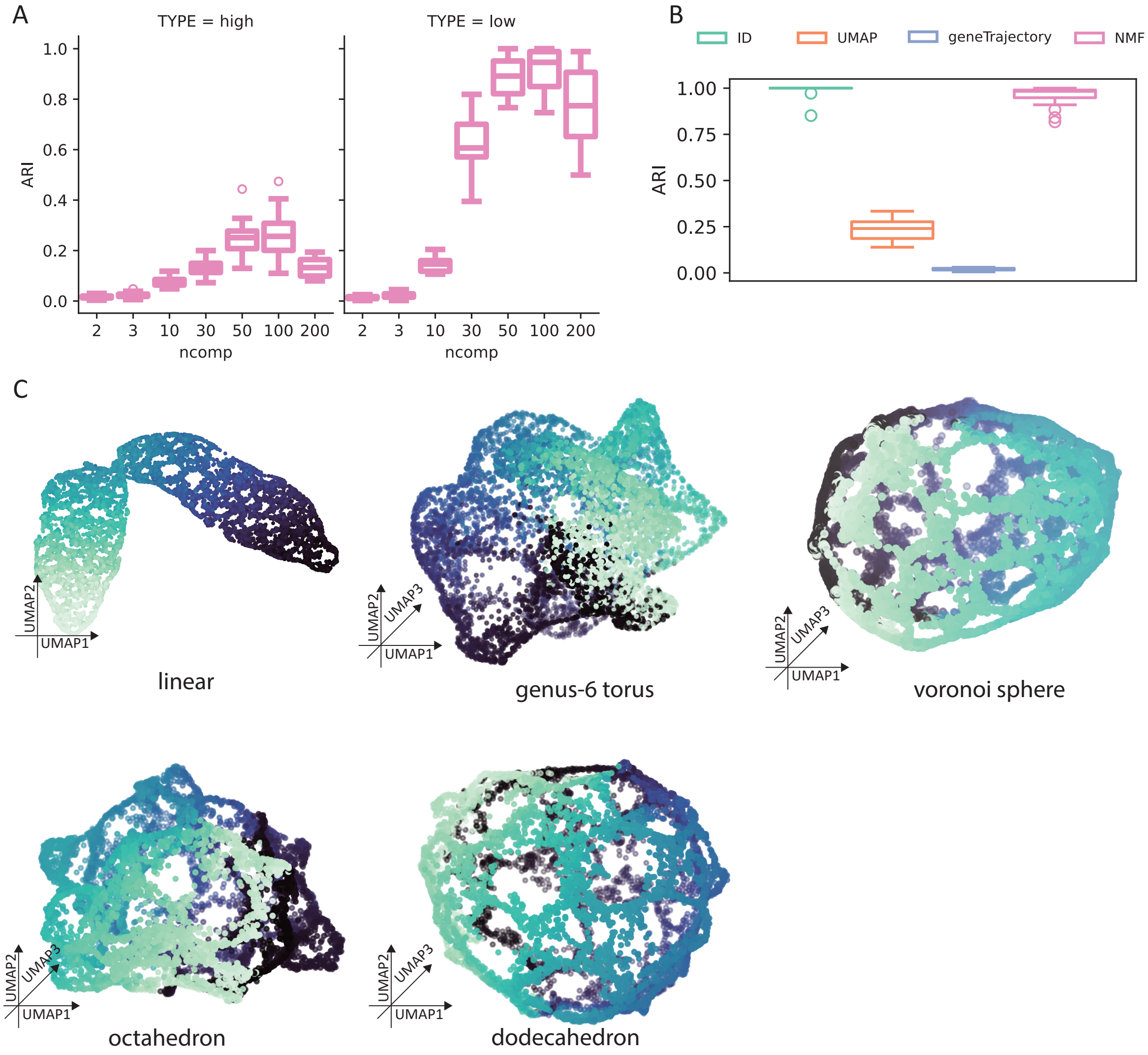


Figure S6: A: ARI computed from the full NMF result (**TYPE=high**) or the full NMF result projected onto 2D using UMAP (**TYPE=low**) with $k$-means clustering over 20 independent trials. B: ARI index computed using k-means using embedding for genes constructed using different methods. 20 individual experiments have been conducted. C: UMAP projection of cells constructed using each cluster of features illustrated in Figure 3C individually.

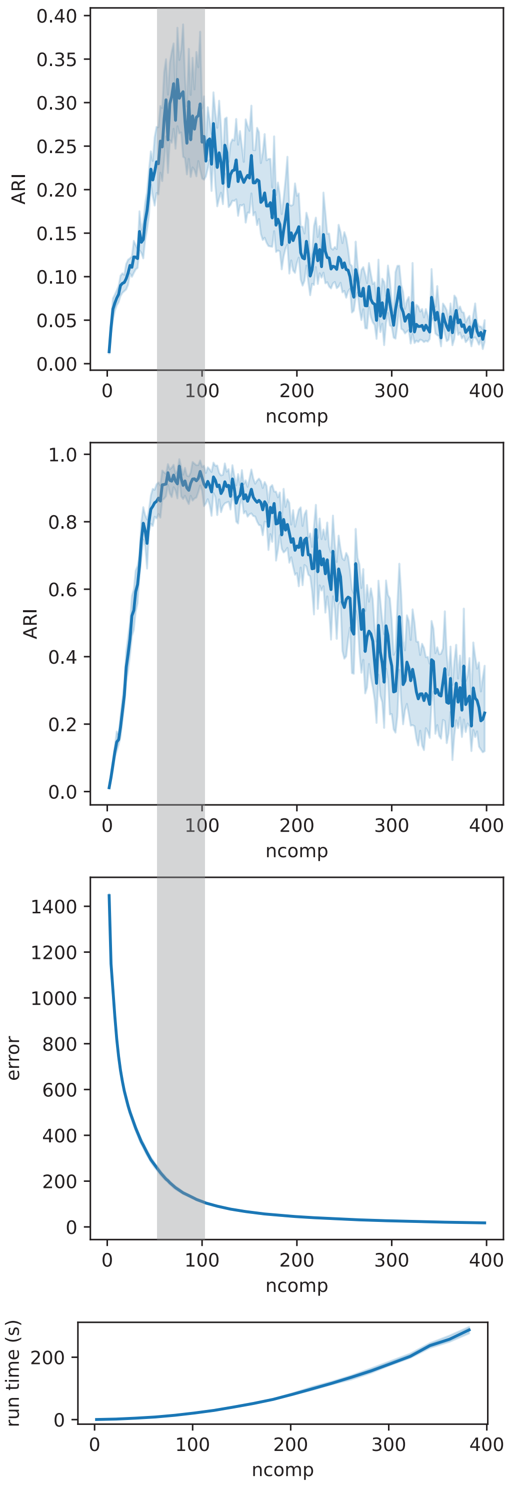


Figure S7: A fine sweep of **ncomp** for NMF using the toy dataset constructed for Figure 3. ARIs were computed using $k$-means clustering in either the high-dimensional space (top) or the low-dimensional space (second) as the number of components (**ncomp**) varies. The third row illustrates how the NMF error changes as the number of components varies. The shaded region denotes the location where the ARI peak occurs when clustering is done in the high-dimensional space. Bottom run time for NMF as a function of the number of components. Each trial was repeated 10 times.

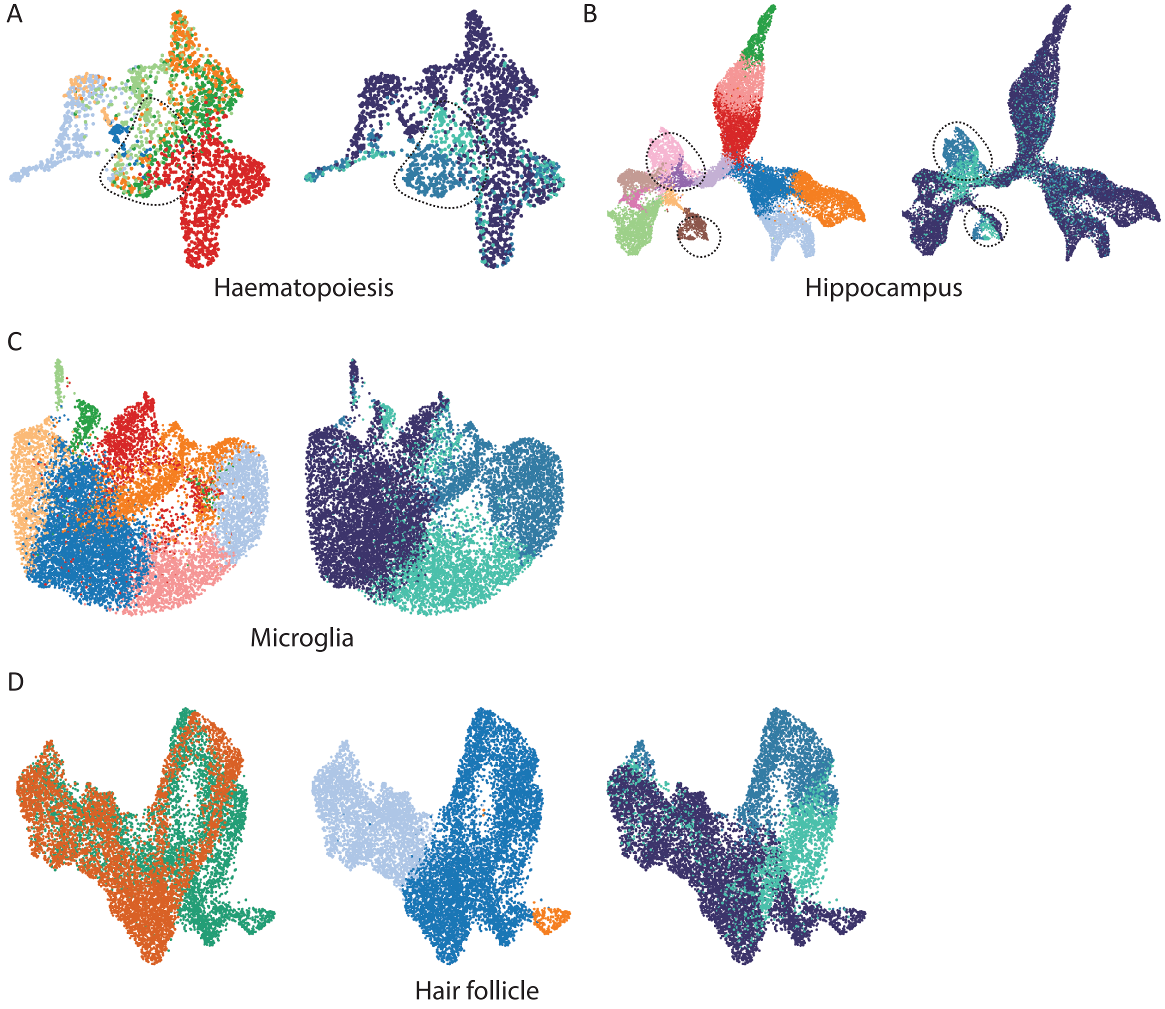


Figure S8: UMAP projection of cells sampled from (A) hematopoetic differentiation [42, 41], (B) hippocampus development [43, 44], (C) microglia [45], and (D) hair follicle [30] using all feature genes. Colors and legends are the same as Figure 4. In panels A and B, the circled region indicates cell–cycle induced branches.

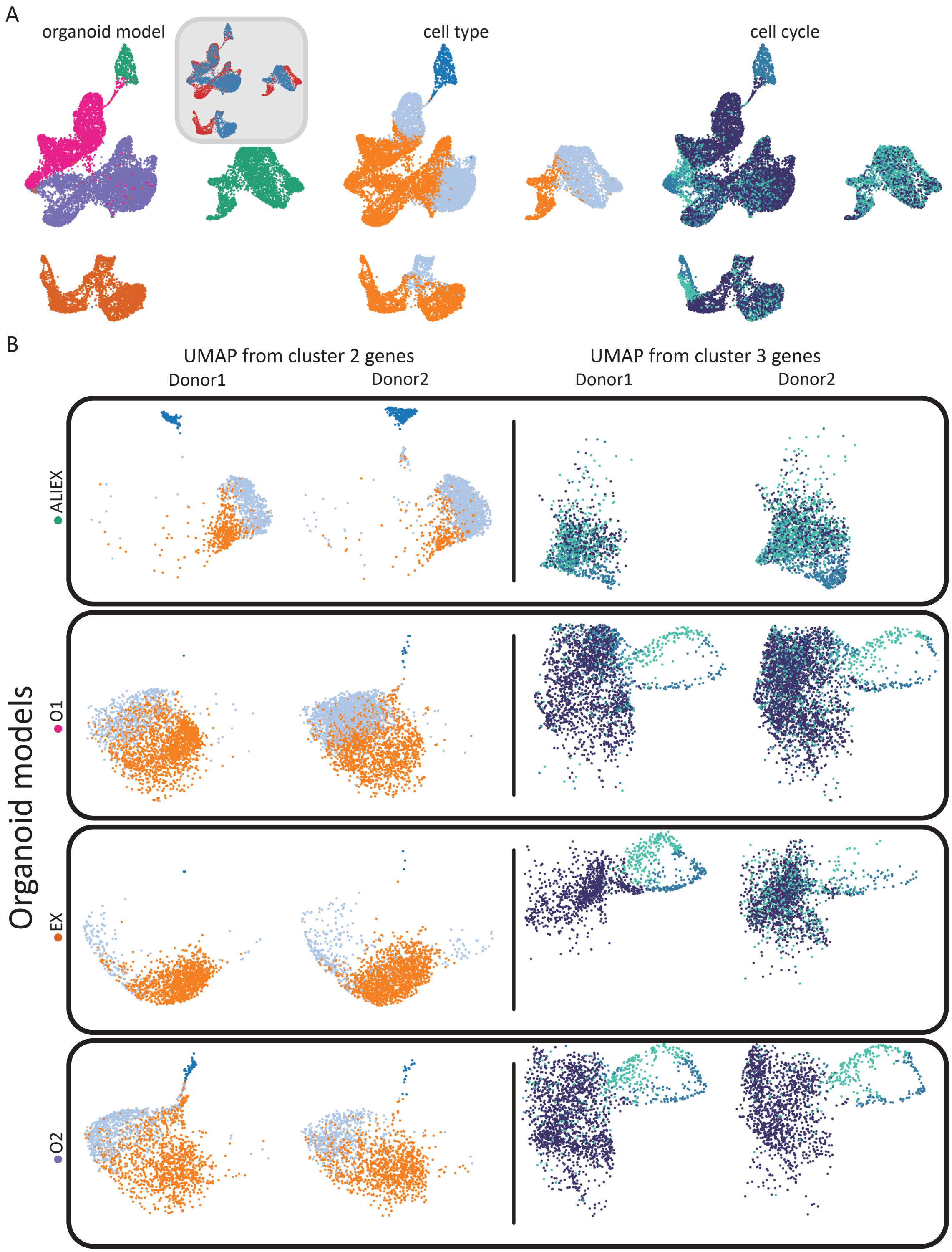


Figure S9: A: UMAP projection of cells from the lung organoid data [55] colored by organoid model, cell type and cell cycle phase (inset: colored by donor). B: UMAP projection of cells using cluster 2 genes only (left) or using cluster 3 genes only (right). Each row indicates a different organoid model, and is subset by donor. Legends are as provided in Figure 5.

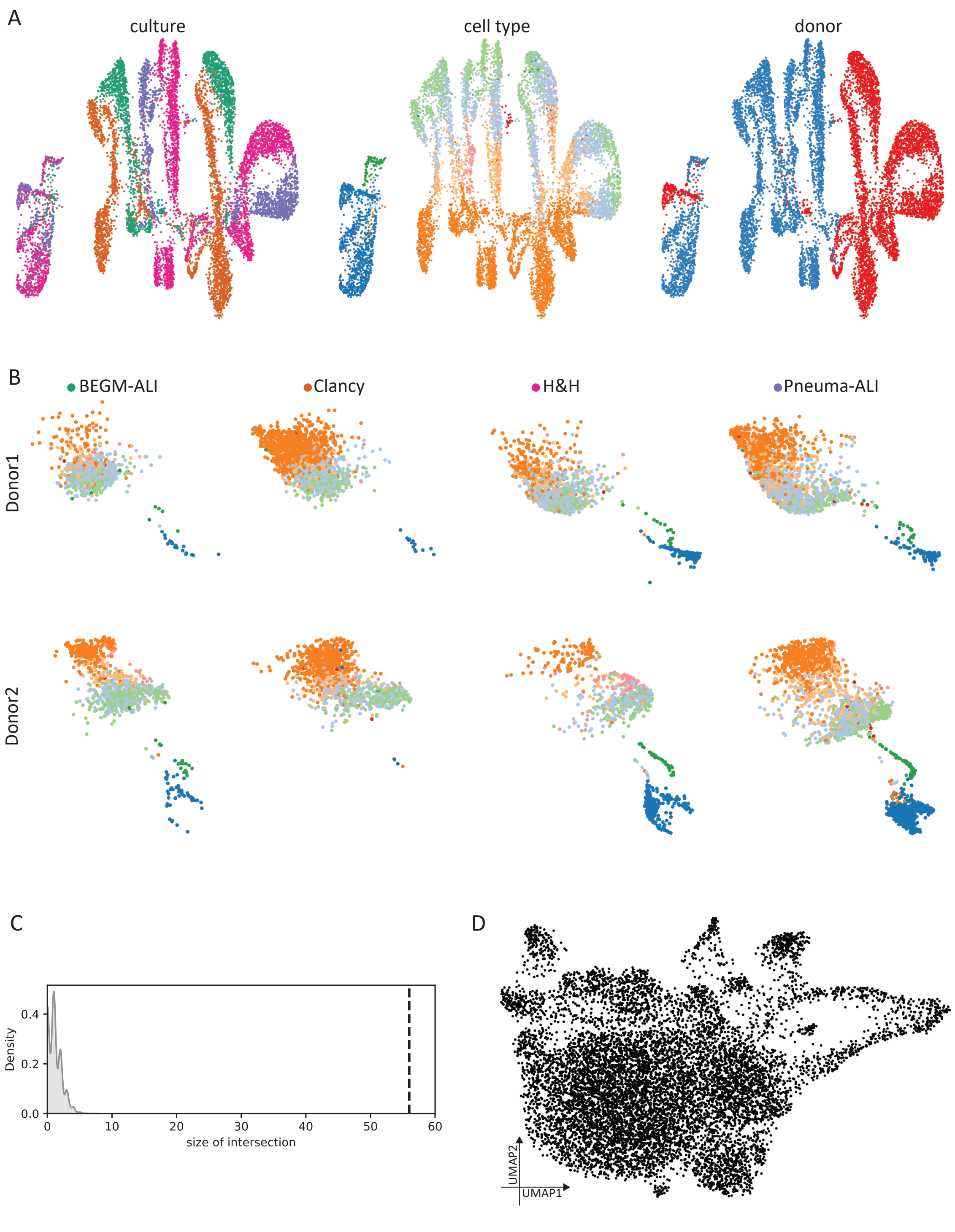


Figure S10: A: UMAP projection of cells from the lung sample data [54] colored by organoid model, cell type and donor. B: UMAP projection of cells using cluster 2 genes only. Each individual panel only contains cells collected from the same donor and cultured with the same media. C: The observed intersection between the cluster 3 genes from both datasets (dashed line) and the null distribution constructed by bootstrapping. D: UMAP projection of cells using cluster 3 genes (from inset of Figure 5A, C) of the lung sample data. Legends are provided in Figure 5.

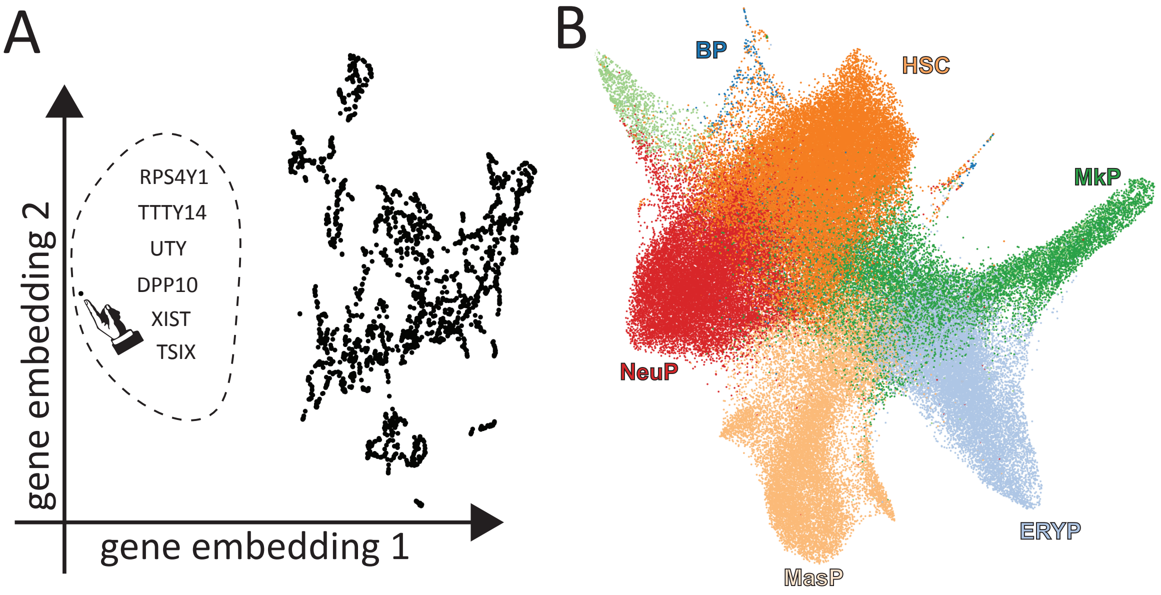


Figure S11: A: Gene embedding of the human HSC dataset; the outlying subject–specific ("batch") cluster is highlighted. B: Low dimensional projection of the data with the six "batch" genes removed, colored by cell type.
